# Keratinocytes coordinate inflammatory responses and regulate development of secondary lymphedema

**DOI:** 10.1101/2023.01.20.524936

**Authors:** Hyeung Ju Park, Raghu P. Kataru, Jinyeon Shin, Gabriela D. García Nores, Elizabeth M. Encarnacion, Mark G. Klang, Elyn Riedel, Michelle Coriddi, Joseph H. Dayan, Babak J. Mehrara

**Affiliations:** Department of Surgery, Division of Plastic and Reconstructive Surgery, Memorial Sloan Kettering Cancer Center, New York, NY

**Keywords:** Secondary lymphedema, Keratinocyte, Hyperkeratosis, Th2-inducing cytokines, PAR2, TSLP, Teriflunomide

## Abstract

Epidermal changes are histological hallmarks of secondary lymphedema, but it is unknown if keratinocytes contribute to its pathophysiology. Using clinical lymphedema specimens and mouse models, we show that keratinocytes play a primary role in lymphedema development by producing T-helper 2 (Th2) -inducing cytokines. Specifically, we find that keratinocyte proliferation and expression of protease-activated receptor 2 (PAR2) are early responses following lymphatic injury and regulate the expression of Th2-inducing cytokines, migration of Langerhans cells, and skin infiltration of Th2-differentiated T cells. Furthermore, inhibition of PAR2 activation with a small molecule inhibitor or the proliferation inhibitor teriflunomide (TF) prevents activation of keratinocytes stimulated with lymphedema fluid. Finally, topical TF is highly effective for decreasing swelling, fibrosis, and inflammation in a preclinical mouse model. Our findings suggest that lymphedema is a chronic inflammatory skin disease, and topically targeting keratinocyte activation may be a clinically effective therapy for this condition.

## Introduction

Lymphedema is a chronic condition caused by inadequate lymphatic function, resulting in cutaneous swelling, fibroadipose deposition, and hyperkeratosis.(*1*) In developed countries, the most common cause of lymphedema is lymph node excision during cancer surgery. It is estimated that 25-40% of patients who undergo surgical treatment for solid tumors develop lymphedema.(*2*) Current treatments for secondary lymphedema—decongestive therapy, compression garments, external pumps, and surgery—are inadequate and costly.(*3, 4*) Likewise, surgical treatments designed to bypass lymphatic channels or promote development of collateral lymphatics are helpful in some patients; however, these procedures are not effective for patients with advanced disease and can cause additional morbidity.(*5*)

Several lines of evidence suggest that the pathophysiology of lymphedema is related to chronic cutaneous T-helper cell inflammatory responses.(*6-14*) CD4^+^ T cell abundance is increased in clinical biopsy specimens, and this inflammatory response positively correlates with severity of disease.(*15*) Depletion of CD4^+^ T cells (but not CD8^+^ cells, natural killer cells, macrophages, or B cells) in mouse models prevents the development of lymphedema and effectively treats established disease.(*6-8, 16, 17*) Topical delivery of tacrolimus, a drug that inhibits T cell proliferation, is highly effective for treating lymphedema in mouse models.(*8*) Recent studies have shown that T-helper 2 (Th2) inflammatory responses and arachidonic acid metabolites play an important role in the pathophysiology of lymphedema by promoting fibrosis and lymphatic leakiness, and impairing pumping of collecting lymphatic.(*3, 8, 13, 15, 16, 18, 19*) Th2 differentiation of naïve CD4^+^ cells is necessary for lymphedema development, since inhibition of this response with neutralizing antibodies targeting IL4 or IL13 or in genetic models deficient in Th2 differentiation is also effective for treating the disease.(*15, 18*) In patients with breast cancer–related lymphedema (BCRL), we found that once monthly infusion of IL4/IL13 neutralizing antibodies significantly improved histologic skin abnormalities and decreased the symptoms of the disease.(*20*) These findings are supported by clinical trials and mouse studies demonstrating that doxycycline is effective for treating filariasis-induced secondary lymphedema by decreasing Th2 immune responses.(*21, 22*) While it is clear that Th2 inflammatory responses are necessary and sufficient for lymphedema development, there remains a significant gap in knowledge on how these responses are activated, which hinders development of new therapies for lymphedema.

Epidermal changes are a prominent finding in lymphedema and include hyperkeratosis, acanthosis, spongiosis, and parakeratosis with elongated rete edges.(*23, 24*) These skin changes are similar to the epidermal changes in atopic dermatitis (AD).(*25*) As with lymphedema, the pathology of AD is also regulated by Th2 inflammatory responses— importantly, epidermal changes in AD are a primary event and precede infiltration of Th2 cells in the skin. Keratinocytes regulate Th2 inflammatory responses and development of AD by producing Th2-inducing cytokines, such as thymic stromal lymphopoietin (TSLP), IL33, and IL25. These cytokines act on naïve CD4^+^ cells through dendritic cells (DCs) to prime Th2 differentiation, regulate cytokine and migratory responses of antigen-presenting cells, and stimulate proliferation of granulocytes that release Th2 cytokines.(*26-33*) The importance of Th2 response in AD is highlighted by the efficacy of dupilumab, a monoclonal antibody that prevents IL4/IL13 signaling.(*34*) Thus, the parallels between AD and lymphedema suggest that keratinocytes may play an important role in the pathophysiology of secondary lymphedema.

In this study, we tested the hypothesis that keratinocytes play a central role in the pathophysiology of lymphedema. Using clinical lymphedema biopsy specimens and mouse models, we show increased expression of protease-activated receptor 2 (PAR2), a sensor for proteolytic enzymes that regulates the expression of Th2-inducing cytokines by keratinocytes in AD,(*35, 36*) and Th2-inducing cytokines by keratinocytes in lymphedema. Furthermore, we show that proliferation and activation of keratinocytes are induced by exposure to lymphatic fluid, and inhibition of PAR2 activation or proliferation attenuate the expression of Th2-inducing cytokines. Finally, we find that inhibition of keratinocyte proliferation decreases the pathology of lymphedema, revealing a potential new target for treating lymphedema.

## Results

### Lymphedema results in hyperkeratosis, de-differentiation of epidermal cells, and increased expression of PAR2 and Th2-inducing cytokines

To analyze epidermal changes resulting from lymphedema, we analyzed skin biopsy samples from the normal and lymphedematous arms of 25 patients with unilateral stage I-II upper extremity BCRL (**Fig. 1A**). RNA sequencing (RNAseq) analysis showed evidence of a Th2 inflammatory response and increased expression of keratins (KRT; **fig. S1A**) in lymphedematous samples, including KRT6 which is indicative of highly proliferative and activated keratinocytes usually observed only in pathological conditions,(*37*) and KRT14 which is expressed by mitotically active, less differentiated keratinocytes typically found in the basal layer of the skin.(*37, 38*) Using qPCR, we confirmed KRT6, KRT14, and KRT16 expression were significantly increased in the lymphedematous versus the normal arm, although there was some inter-patient variability (**Fig. 1B**). This variability was not related to the severity or duration of the disease (Data not shown). Histological and protein analysis of skin biopsy samples showed that lymphedema was associated with hyperkeratosis, increased epidermal area, increased number of proliferating Ki67^+^ keratinocytes, and increased expression of KRT6 and KRT14 (**Fig. 1, C to E**). In addition, KRT6-expressing keratinocytes were enlarged and abnormal in appearance, while KRT14 expression was noted in all layers of the epidermis of lymphedematous arms, indicating decreased differentiation (**Fig. 1C**). These keratinocyte changes correlated with higher expression of keratinocyte growth factors (EGF, EGFR, IL1α) in lymphedematous versus normal skin (**fig. S1B**).

**Figure 1.**
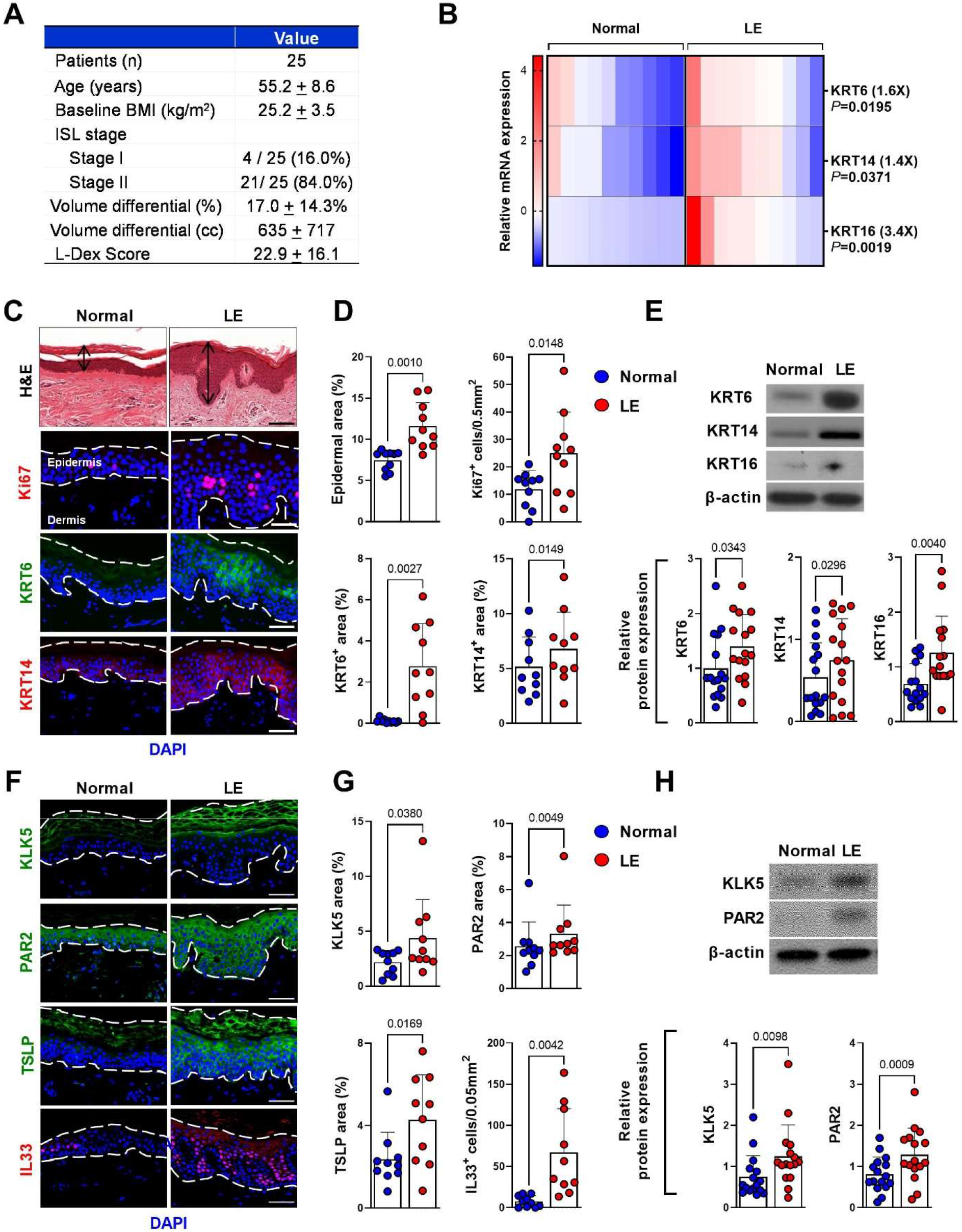
Lymphedema results in hyperkeratosis and expression of Th2-inducing cytokines. A. Demographics of patients with unilateral BCRL who provided samples for our study. Data are mean ± standard deviation unless noted. BMI: body mass index; ISL: International Society of Lymphology. B. Relative mRNA expression by qPCR of KRT6, KRT14, and KRT16 in normal and lymphedematous (LE) skin biopsies from patients with unilateral BCRL (N=10). mRNA expression was normalized to β-actin expression. Each box is representative of one patient. *P* values were calculated by paired student’s t-test. Fold change comparing LE versus normal skin biopsy is shown in parentheses. C. Representative images of H&E and immunofluorescent staining of Ki67, KRT6, and KRT14 in normal and lymphedematous (LE) skin biopsies from patients with unilateral BCRL. Arrow indicates epidermis. Dashed lines indicate the thickness of epidermis. Scale bars: 100 µm (H&E), 50 µm (IHC). D. Quantification of epidermal area, Ki67^+^ cells, and KRT6 and KRT14 area in normal and lymphedematous (LE) skin biopsies from patients with unilateral BCRL. Each circle represents the average quantification of 3 high-power field (HPF) views for each patient (N=10) *P* values were calculated by paired student’s t-test. E. Representative western blot images (*top*) and quantification (*bottom*; relative to β-actin) of KRT6, KRT14, and KRT16 in normal and lymphedematous (LE) skin biopsies from patients with unilateral BCRL. Each circle represents one patient (N=16). *P* values were calculated by paired student’s t-test. F. Representative immunofluorescent images of KLK5, PAR2, TSLP, and IL33 staining in normal and lymphedematous (LE) skin biopsies from patients with unilateral BCRL. Dashed lines indicate the thickness of epidermis. Scale bar: 50 µm. G. Quantification of KLK5, PAR2, TSLP, and IL33 area in normal and lymphedematous (LE) skin biopsies from patients with unilateral BCRL. Each circle represents the average quantification of 3 HPF views for each patient (N=10). *P* values were calculated by student’s t-test. H. Representative western blots (*top*) and quantification (*bottom*; relative to β-actin) of KLK5 and PAR2 in normal and lymphedematous (LE) skin biopsies from patients with unilateral BCRL. Each circle represents one patient (N=16). *P* values were calculated by paired student’s t-test.

Previously PAR2 was shown as a regulator of Th2-inducing cytokines in AD. PAR2 is activated by serine proteases, such as Kallikrein 5 (KLK5), which cleave N-terminal of the PAR2 molecule to expose the tethered ligand. We found the expression of KLK5, PAR2, TSLP, and IL33 markedly increased in lymphedematous compared with normal skin (**Fig, 1, F to H**). KLK5 staining was localized to the cornified layer of the skin, while PAR2, TSLP, and IL33 staining were present and increased in the entire epidermis (**Fig, 1F**). We used negative (no primary antibody) controls to confirm the specificity of our findings (**fig S1C**). PAR2 activates the expression of Th2-inducing cytokines by NFATc1 activation,(*39*) we consistently found a significant increase in NFATc1 staining in lymphedematous versus normal skin (**fig S1B**).

### Keratinocyte expression of Th2-inducing cytokines occurs rapidly after lymphatic injury and precedes CD4^+^ cell inflammatory responses

We next used a mouse tail model of lymphedema to understand the temporal changes in the epidermis relative to the timing of lymphatic injury and the development of lymphedema. In this model, histological signs of lymphedema, such as inflammation and fibroadipose deposition, develop 4-6 weeks after skin and lymphatic excision.(*16, 40*) We therefore harvested tail skin specimens 2 and 6 weeks after surgery to analyze epidermal changes before and after the onset of lymphedema.(*40*) We found that hyperkeratosis occurred rapidly after lymphatic injury and was evident even at the 2-week time point in skin sections harvested 2-3 cm from the excision site (**Fig. 2A and B**). Hyperkeratosis increased significantly by 6 weeks after surgery suggesting that epidermal changes, similar to lymphedema, are progressive in nature. Importantly, we found that epidermal changes preceded dermal infiltration of CD3^+^ T cells suggesting that epidermal Th2-inducing cytokine expression initiates the pathology of lymphedema (**Fig. 2A and B**).

**Figure 2.**
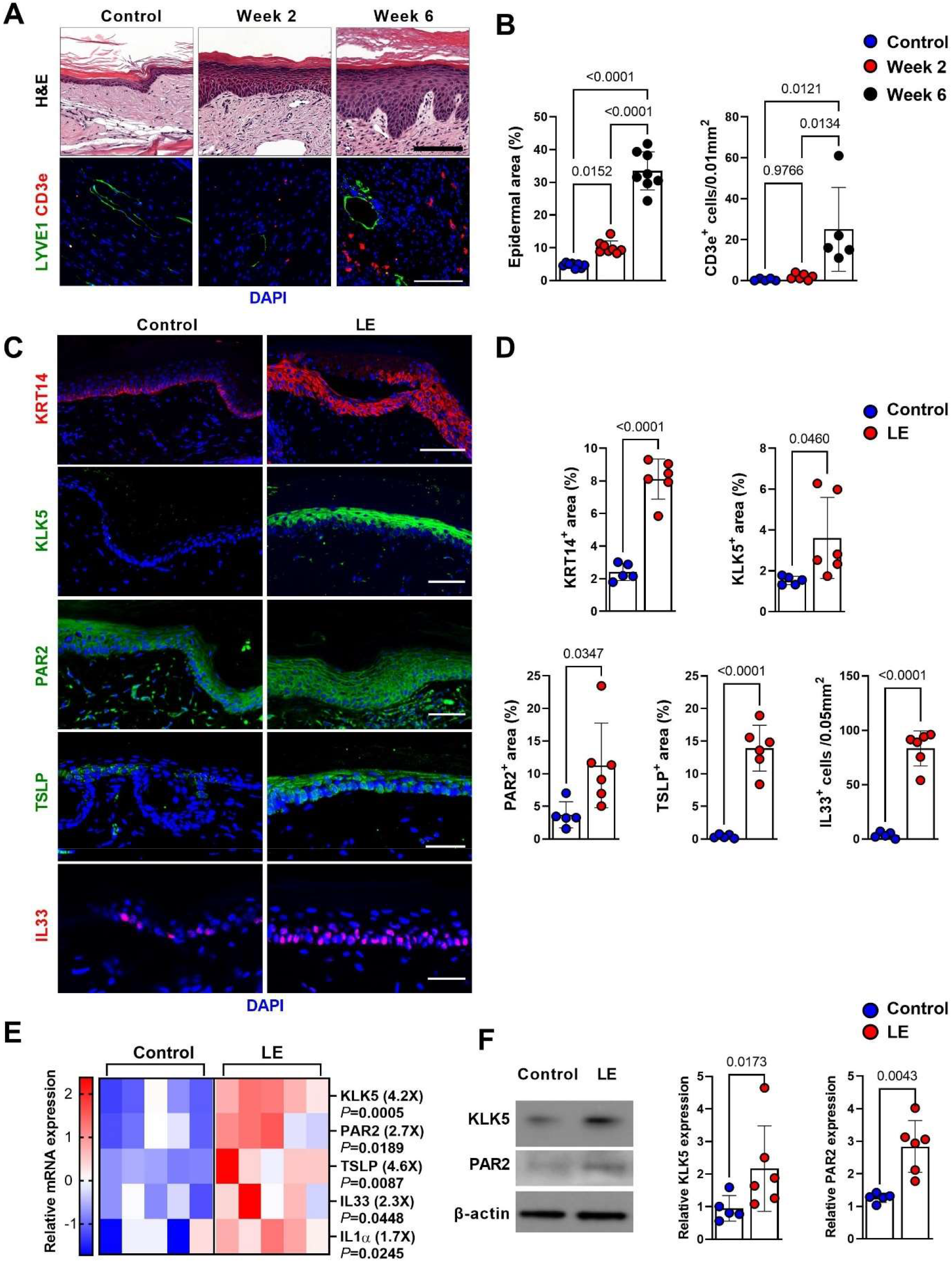
Keratinocyte expression of Th2-inducing cytokines occurs rapidly after lymphatic injury. A. Representative H&E (*top*) and immunofluorescent staining for LYVE1 and CD3e (*bottom*) of control (skin incision) tail skin and tail skin harvested 2 or 6 weeks after tail skin and lymphatic excision. Scale bar: 100 µm. B. Quantification of epidermal area and CD3e^+^ cells in control and tail skin harvested 2 or 6 weeks after tail skin and lymphatic excision. Each circle represents the average of 3 HPF views of each mouse (N=5-8). *P* values were calculated by one-way ANOVA. C. Representative immunofluorescent images of KRT14, KLK5, PAR2, TSLP, and IL33 staining in tail skin harvested 2 weeks after surgery from control and lymphedema (LE) mice. Scale bar: 100 µm. D. Quantification of KRT14, KLK5, PAR2, TSLP, and IL33 area tail skin harvested 2 weeks after surgery from control and lymphedema (LE) mice. Each circle represents the average of 3 HPF views of each mouse (N=5-6). *P* values were calculated by unpaired student’s t-test. E. Relative mRNA expression by qPCR tail skin harvested 2 weeks after surgery from control and lymphedema (LE) mice (N=5). mRNA expression was normalized to β-actin expression. Each box represents one mouse. *P* values were calculated by Mann-Whitney test. Fold changes from control are shown in parentheses. F. Representative western blot (*left*) and quantification (*right;* relative to β-actin) of KLK5 and PAR2 in tail skin harvested 2 weeks after surgery from control and lymphedema (LE) mice (*left*). Each circle represents each mouse (N=5-6). *P* values were calculated by Mann-Whitney test.

To control for the effects of surgery, we next compared tail skin harvested 2 weeks after surgery from mice treated with skin incision alone (control) versus mice that underwent skin/lymphatic excision (lymphedema). We found that the expression of KRT6, KRT14, Ki67, IL1α, KLK5, PAR2, NFATc1, TSLP, and IL33 were increased in mouse tail skin from lymphedema versus control mice before the onset of lymphedema (**Fig. 2, C and D, fig. S2, A and B**). Spatial patterns of these molecules were highly concordant with our skin biopsy samples (**Fig. 1, C and D, fig. S1B**). We confirmed our histological findings with qPCR and found that the expression of KLK5 (4.2-fold), PAR2 (2.7-fold), TSLP (4.6-fold), IL33 (2.3-fold), and IL1a (1.7-fold) were significantly increased in skin specimens from lymphedema compared with control mice (**Figure 2E**). These mRNA changes translated to increased protein expression of KLK5, PAR2, TSLP, and IL33 in lymphedema versus control mice (**Fig. 2F, fig. S2C**).

We and others have shown that CD4^+^ cells play a key role in the pathophysiology of lymphedema.(*6, 15, 16*) Therefore, to determine if epidermal changes are independent of CD4^+^ inflammatory responses, we compared tail skin specimens from wild-type and CD4 knockout mice (CD4KO) after tail skin/lymphatic excision. Consistent with our previous reports, CD4KO mice had decreased tail swelling and evidence of lymphedema compared with wild-type controls (Data not shown).(*16, 18*) However, we found no differences in expression of KRT6, KRT14, KLK5, and PAR2 2 weeks after surgery (**fig. S2, D and E**), suggesting that epidermal changes are independent of T cell inflammatory responses in lymphedema.

The mouse tail model of lymphedema is a surgical procedure, and changes in the epidermis may also reflect wound healing, not just lymphatic injury. We therefore used a non-surgical model of lymphedema using diphtheria toxin (DT) injections in the hindlimb of transgenic mice that express the human diphtheria toxin receptor under the regulation of the Flt4 (Vegfr3) promoter (DTR mice).(*41*) In this model, injection of DT ablates the superficial and deep lymphatics and results in the development of chronic and progressive lymphedema that is histologically and radiologically identical to clinical lymphedema.(*41*) Chronic lymphedema changes, including swelling and T cell infiltration, develop 6-9 weeks after DT injection in this model. Consistent with our findings using the mouse tail model of lymphedema, we found that hyperkeratosis, epidermal proliferation (Ki67^+^ keratinocytes), and expression of KRT6 were increased in the hindlimb skin as early as 3 weeks following DT injection in DTR mice versus wild-type controls (**fig S3, A and B**). These changes were even more pronounced by 9 weeks after DT injection when lymphedema pathology was evident as previously shown.(*41*) Similarly, keratinocyte expression of KLK5, PAR2, TSLP, and IL33 were increased in specimens collected 3 weeks after DT injection and further increased at 9 weeks compared to controls (**fig S3, A and B**). Gene expression confirmed these findings, showing that lymphatic injury markedly increased the expression of KLK5, PAR2, TSLP, IL33, and IL1*α* in FLT4CreDTRfloxed mice (**fig S3C**).

### PAR2 deficiency reduces Th2 inflammation and lymphedema

PAR2 regulates the expression of Th2-inducing cytokines and Th2 inflammatory responses in skin disorders, and inhibition of PAR2 decreases the severity of skin diseases such as AD and Netherton Syndrome.(*42, 43*) To investigate the role of PAR2 of in lymphedema development, we compared swelling and secondary changes of lymphedema (inflammation, fibrosis, and lymphatic dilatation) in wild-type and PAR2 knockout (PAR2KO) mice 6 weeks after tail skin and lymphatic excision. We noted an increase in tail volume in both wild-type and PAR2KO mice early after surgery (1-3 weeks; **Fig. 3B**). However, in contrast to wild-type mice, tail swelling in PAR2KO mice decreased significantly thereafter, and 6 weeks following surgery, PAR2KO mice had a 2-fold decrease in tail swelling that was noticeable even on gross examination (**Fig. 3, A and B**). Loss of PAR2 expression significantly decreased type I collagen deposition and CD4^+^ cell infiltration 6 weeks after surgery (**Fig. 3C, fig. S4A**). Consistent with a decreased lymphedema phenotype, PAR2KO mice also had an increased number of lymphatic vessels and decreased lymphatic vessel diameter compared with wild-type controls (**Fig. 3C, fig. S4A**).

**Figure 3.**
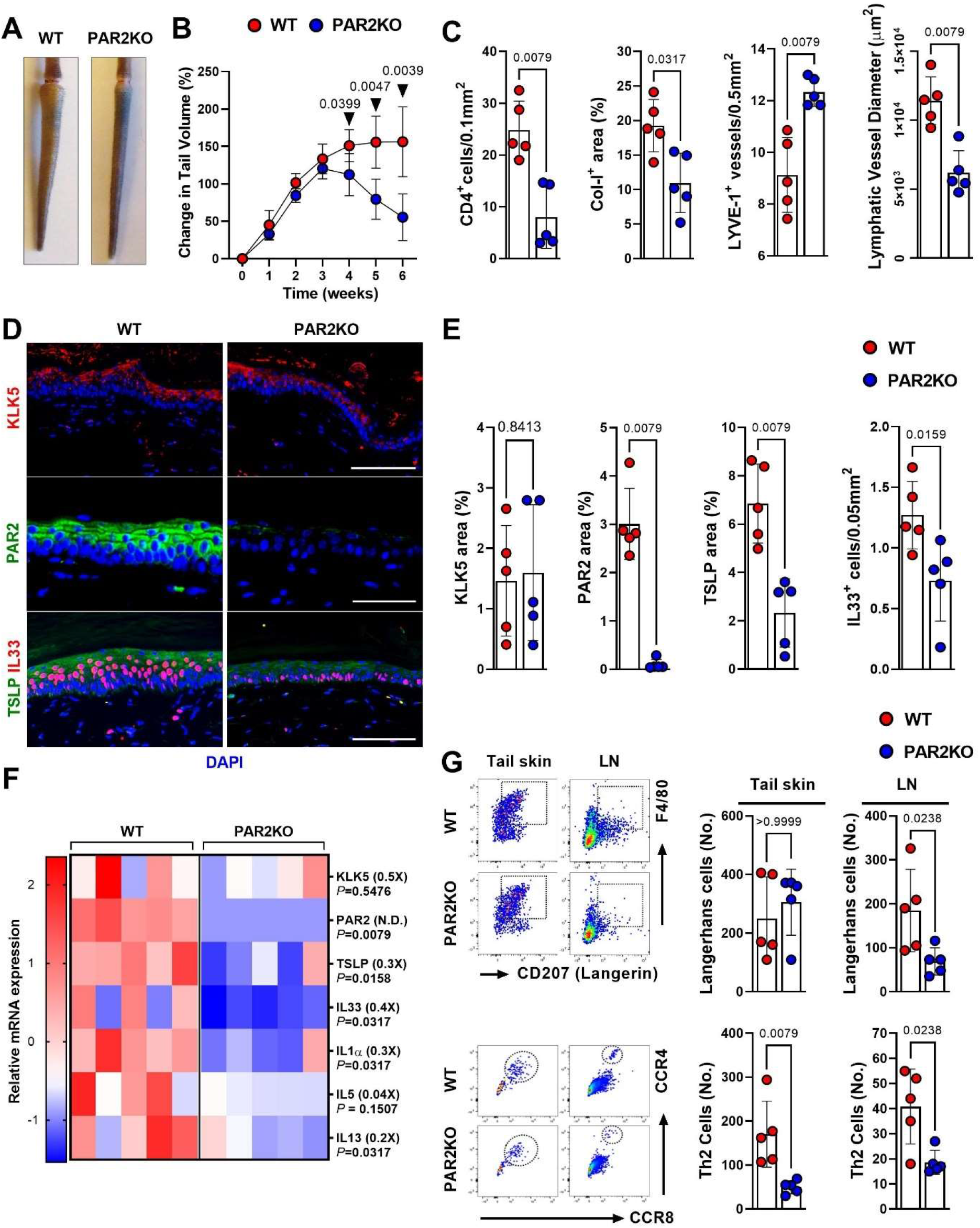
PAR2 knockout reduces lymphedema. A. Representative gross images of tails from wild-type (WT) and PAR2 knockout (PAR2KO) mice harvested 6 weeks after tail skin and lymphatic excision. B. Changes in tail volume over time in WT and PAR2KO mice. Each circle represents the average measurement from each mouse (N=5). *P* values were calculated by two-way ANOVA. C. Quantification of CD4^+^ cells, collagen I, LYVE1^+^ vessels, and lymphatic vessel diameter in tail skin of WT and PAR2KO mice harvested 6 weeks after tail skin and lymphatic excision. Each circle represents the average quantification of 3 HPF views for each mouse (N=5). *P* values were calculated by Mann-Whitney test. D. Representative immunofluorescent images of KLK5, PAR2, TSLP, and IL33 staining in tail skin of WT and PAR2KO mice harvested 6 weeks after tail skin and lymphatic excision. Scale bar: 100 µm. E. Quantification of KLK5, PAR2, TSLP, and IL33 area in tail skin of WT and PAR2KO mice harvested 6 weeks after tail skin and lymphatic excision. Each circle represents the average quantification of 3 HPF views for each mouse (N=5). *P* values were calculated by Mann-Whitney test. F. Relative mRNA expression by qPCR in tail skin of WT and PAR2KO mice harvested 6 weeks after tail skin and lymphatic excision (N=5). mRNA expression was normalized by β-actin expression. Each box represents one mouse. *P* values were calculated by Mann-Whitney test. Fold change from control is shown in parentheses. G. Representative flow cytometry of Langerhans cells and Th2 cells from tail skin and draining lymph nodes (LN) of WT and PAR2KO mice harvested 6 weeks after tail skin and lymphatic excision (*left*). Quantification of the number of Langerhans cells and Th2 cells (*right*). Each circle represents each mouse (N=5). *P* values were calculated by Mann-Whitney test.

Loss of PAR2 also decreased hyperkeratosis and expression of KRT6 and Ki67 relative to controls **(fig. S4B)**. These skin changes correlated with decreased keratinocyte expression of TSLP, IL33, KRT6, and NFATc1, but not KLK5 which is upstream of PAR2 (**Fig. 3, D to F, fig. S4, B and C**). We also examined the number of Langerhans cells (LCs) and Th2 cells in skin and draining lymph nodes 6 weeks after tail skin and lymphatic excision and found that loss of PAR2 did not significantly alter the number of activated LCs in the skin but decreased the number of LCs in the draining lymph nodes. PAR2KO mice also had significantly decreased numbers of Th2 cells infiltrating the skin and draining lymph nodes (**Fig. 3G**).

### Lymphatic fluid activates keratinocyte proliferation and cytokine expression by a PAR2-dependent mechanism

To investigate how lymphedema induces skin changes, we cultured human keratinocytes (h-keratinocytes) with or without lymphedema fluid (LF; from patients with stage II BCRL) and found that LF significantly increased h-keratinocyte expression of KRT6, Ki67, KLK5, PAR2, and TSLP (**Fig. 4, A and B**). The addition of a small molecule inhibitor of PAR2 (ENMD1068) decreased expression of KRT6 and PAR2 to control, and significantly decreased expression of Ki67, KLK5, and TSLP compared to LF alone (**Fig. 4, A and B**). mRNA expression confirmed that exposure of keratinocytes to LF markedly increased the expression of these factors and that these changes were abrogated by EMND1068 (**fig. S5A**).

**Figure 4.**
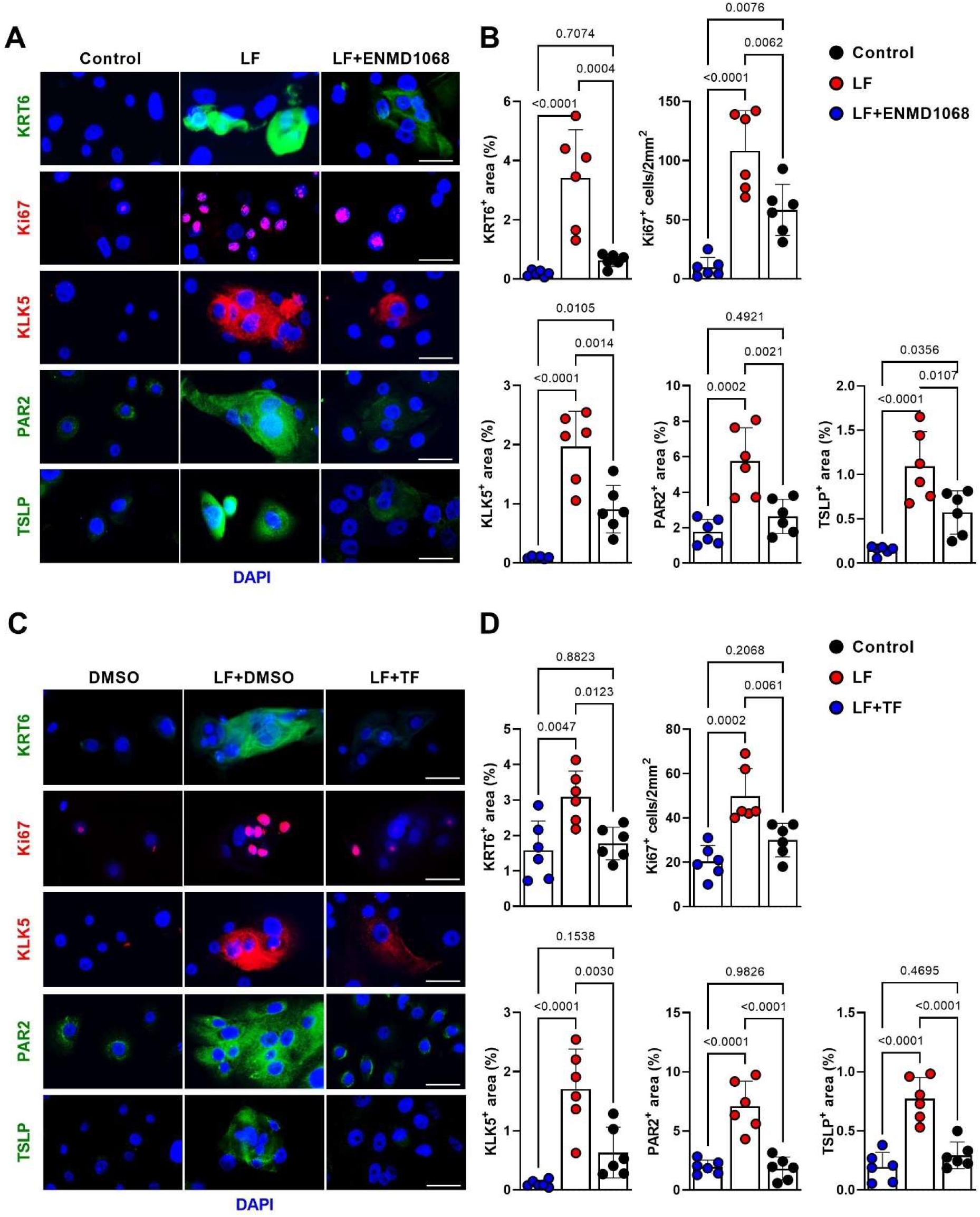
Lymphedema fluid induces proliferation and increases expression of PAR2 and Th2-inducing cytokines in human keratinocytes. A. Representative immunofluorescent images of KRT6, Ki67, KLK5, PAR2, and TSLP staining in h-keratinocytes cultured with PBS (control), lymphatic fluid (LF), or LF+ENMD1068. Scale bar: 20 µm. B. Quantification of KRT6, Ki67, KLK5, PAR2, and TSLP area in h-keratinocytes cultured with PBS (control), LF, or LF+ENMD1068. Each circle represents the average quantification of 2 HPF views for each experiment. Lymph fluid from 2 different lymphedema patients was used (N=6; 3 for each lymph fluid). *P* values were calculated by one-way ANOVA. C. Representative immunofluorescent images of KRT6, Ki67, KLK5, PAR2, and TSLP staining of h-keratinocytes cultured with DMSO, LF+DMSO, or LF+TF (25 µM in DMSO). Scale bar: 20 µm. D. Quantification of KRT6, Ki67, KLK5, PAR2, and TSLP area in h-keratinocytes cultured with DMSO, LF, or LF+TF. Each circle represents the average quantification of 2 HPF views for each experiment (N=6). *P* values were calculated by one-way ANOVA.

As LF caused keratinocyte proliferation, we next tested if inhibition of this response could decrease PAR2 and Th2-inducing cytokine expression. Teriflunomide (TF), the active metabolite of leflunomide, is an FDA-approved treatment for multiple sclerosis.(*44*) TF inhibits de novo synthesis of pyrimidine by blocking dihydroorotate dehydrogenase thus inhibiting cellular proliferation.(*45*) Consistently, TF significantly abrogated h-keratinocyte proliferation *in vitro* (**fig. S5B**). Furthermore, TF (25 µM) significantly decreased the expression of KRT6, Ki67, KLK5, PAR2, and TSLP in LF-stimulated h-keratinocytes, similar to our findings with ENMD1068 (**Fig. 4, C and D**). In fact, TF was even more effective than ENDM1068 in decreasing TSLP expression to baseline levels.

### TF decreases epidermal changes and other pathological changes of secondary lymphedema

Since TF was highly effective in preventing pathological changes of h-keratinocytes to LF, we next tested topical TF *in vivo* in our lymphedema mouse model. We used Aquaphor® as a carrier for TF based on our prior experience with topical formulations for lymphedema.(*8, 40*) Two weeks after tail skin/lymphatic excision surgery, control mice were treated with Aquaphor® ointment applied to the distal tail skin, and experimental animals were treated with Aquaphor® containing TF (27 mg/ml) once a day for 4 weeks. Within 1 week of treatment, mice treated with topical TF had decreased swelling and tail volumes compared with controls (**Fig. 5, A and B**). Improvements in lymphedema in TF-treated mice continued, such that 4 weeks after initiation of treatment, TF-treated mice had virtually no swelling. Consistent with improved lymphedema outcomes, we also found that mice treated with topical TF had decreased dermal infiltration of CD4^+^ cells, decreased type I collagen deposition, increased number of lymphatic capillaries, decreased lymphatic vessel diameter, and decreased expression of KLK5, PAR2, TSLP, and IL33 compared to control mice (**Fig. 5C, fig. S6, A to C**). To investigate if treatment with topical TF can mitigate early pathological changes in keratinocytes following lymphatic injury, we repeated the experiment, instead beginning TF treatment immediately after surgery and dosing once daily for 2 weeks. This approach reduced tail swelling and markedly decreased skin expression of KLK5, PAR2, TSLP, IL33, Ki67, KRT6, IL1*α*, and NFATc1 (**Fig. 5, D to G, fig. S6, D to F**).

**Figure 5.**
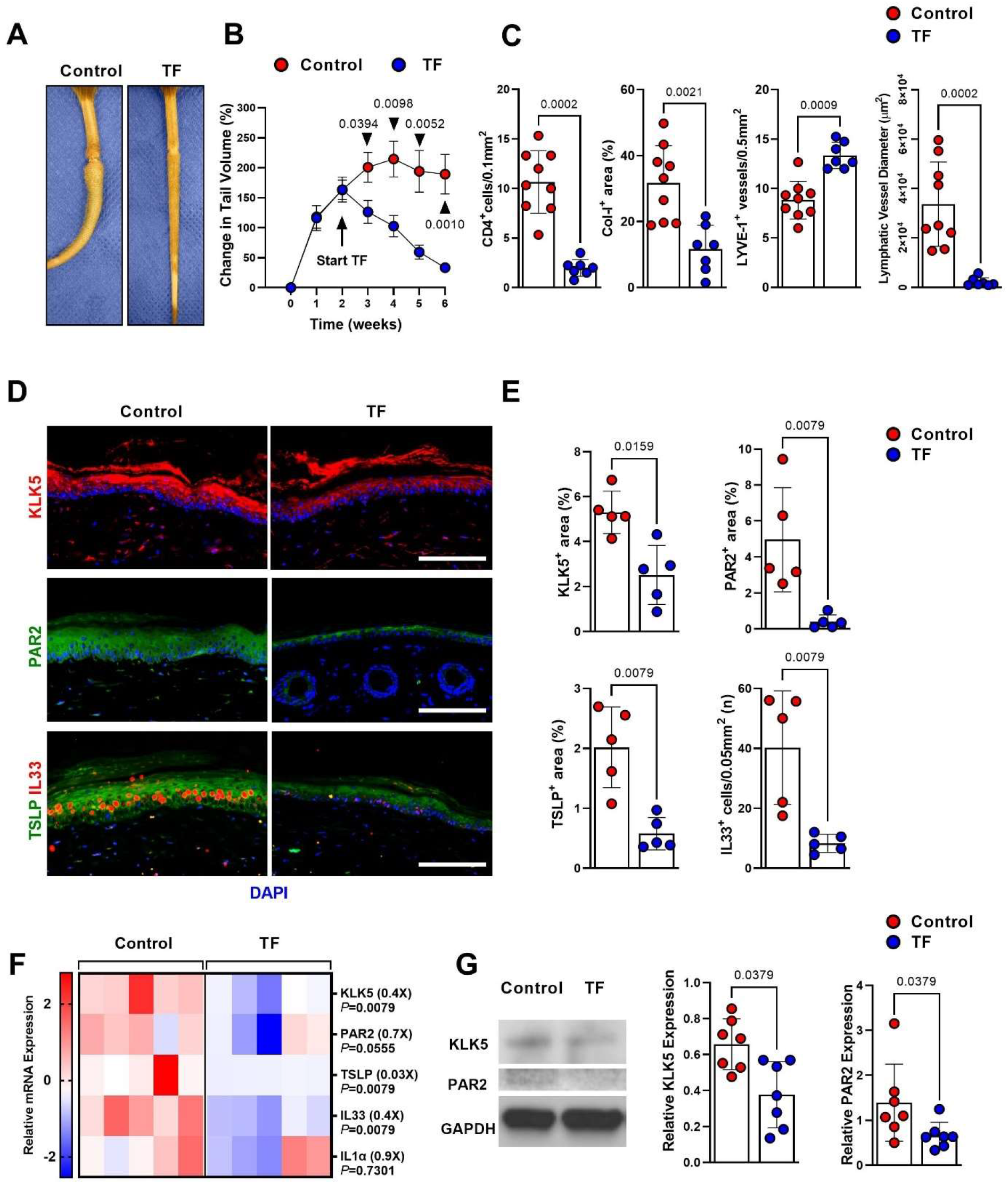
Topical treatment with teriflunomide prevents lymphedema development. A. Representative images of control and teriflunomide (TF) -treated mice 6 weeks after tail skin and lymphatic excision. B. Changes in tail volume over time in mice treated with vehicle (control) or TF once daily for 4 weeks beginning 2 weeks after tail skin and lymphatic excision. Each circle represents the average measurement from each mouse (N=7-9). *P* values were calculated by two-way ANOVA. C. Quantification of CD4^+^ cells, collagen I, LYVE1^+^ vessels, and lymphatic vessel diameter in skin samples harvested from mice treated with vehicle (control) or TF once daily for 4 weeks beginning 2 weeks after tail skin and lymphatic excision. Each circle represents the average quantification of 3 HPF views for each mouse (N=7-9). *P* values were calculated by Mann-Whitney test. D. Representative immunofluorescent images of KLK5, PAR2, TSLP, and IL33 staining in mouse tails from mice treated with vehicle (control) or TF for 2 weeks starting one day after tail skin and lymphatic excision. Scale bar: 100 µm. E. Quantification of KLK5, PAR2, TSLP, and IL33 area in tail skin from mice treated with vehicle (control) or TF for 2 weeks starting one day after tail skin and lymphatic excision. Each circle represents the average quantification of 3 HPF views for each mouse (N=5). *P* values were calculated by Mann-Whitney test. F. Relative mRNA expression by qPCR in tail skin from mice treated with vehicle (control) or TF for 2 weeks starting one day after tail skin and lymphatic excision (N=5). mRNA expression was normalized to β-actin expression. Each box represents one mouse. *P* values were calculated by Mann-Whitney test. Fold change from control is shown in parentheses. G. Representative images of western blots (*left*) and quantification (*right*; relative to GAPDH) of KLK5 and PAR2 in tail skin from mice treated with vehicle (control) or TF. Each dot represents quantification of a separate Western blot (N=7). *P* values were calculated by Mann-Whitney test.

To ensure the observed improvements following topical TF treatment were not simply due to improvements in surgical wound healing, we next analyzed the efficacy of TF treatment in our non-surgical model of lymphedema. A week after the last DT injection, control mice were treated with Aquaphor®, while experimental mice were treated with Aquaphor® containing TF once daily for 8 weeks. Consistent with the tail model, gross analysis of the hindlimbs revealed significant swelling in the distal hindlimb of control mice; in contrast, TF-treated mice had minimal swelling (**fig. S7A**). Mice treated with topical TF also had markedly decreased hyperkeratosis, decreased expression of KRT6, decreased number of Ki67^+^ cells in the epidermis, and decreased expression of KLK5, PAR2, TSLP, IL33, and IL1*α* (**fig. S7, A to D**).

## Discussion

The skin is the largest organ in the body and consists of the epidermis, dermis, and hypodermis. Keratinocytes are skin cells that that make up 90% of the epidermis and originate as stem cells in basal layers of the skin. Keratinocytes proliferate, differentiate, and migrate to the more superficial layers of the skin ultimately forming the cornified layer. Major functions of keratinocytes include maintenance of skin barrier function, prevention of water loss, and inhibition of bacterial infiltration.(*46*) Proliferation and differentiation of keratinocytes is controlled by diverse cytokines such as Interleukin (IL) 1α, IL1β, epithelial growth factor (EGF), transforming growth factor 1α (TGF1α) and tumor necrosis factor α (TNFα).(*47, 48*) Keratins (KRT) are a large family of intermediate filaments that are expressed by keratinocytes and are necessary for maintenance of cytoskeletal integrity and cellular motility.(*37*) KRT14-KRT5 are expressed by keratinocytes and keratinocyte precursors in the stratum basale. As the cells migrate suprabasally and become differentiated, expression of KRT14-KRT5 heterodimers is replaced by KRT10-KRT1. KRT16, KRT17, and KRT6 are expressed in activated, proliferating keratinocytes in pathological or physiological conditions such as atopic dermatitis, psoriasis, wound healing, and burns.

Hyperkeratosis is a histological hallmark of lymphedema and is a common finding in inflammatory skin disorders.(*23, 24*) Although keratinocytes are known to play a key role in the pathophysiology of psoriasis and AD, no prior studies have tested the hypothesis that these cells also contribute to the pathology of secondary lymphedema. In this study, we used clinical samples collected from women with unilateral BCRL to show that lymphedema increases proliferation and decreases differentiation of keratinocytes in the basal layer of the skin.

Keratinocytes in lymphedematous skin also exhibit stress and activation, as evidenced by increased expression of KRT6, KRT16, and KRT17. Using mouse models of lymphedema, we found that epidermal changes occur rapidly after lymphatic injury and that these changes precede infiltration of CD4^+^ cells. We also found that lymphedema increases the expression of EGF, EGFR, and IL1*α*. Importantly, we found that both surgical and non-surgical models of lymphedema have the same epidermal changes, suggesting this is independent of wound healing. LF activated h-keratinocyte proliferation and expression of KRT6 in vitro, suggesting that soluble factors in lymphedematous tissues regulate keratinocyte changes. These findings are consistent with a previous report demonstrating that LF induced keratinocyte proliferation and expression of KRT6, KRT16, and KRT17 and that this response was mitigated by inhibiting IL1β, keratinocyte growth factor, or TNF-*α*.(*23*) Taken together, our data suggest that lymphatic injury rapidly activates keratinocytes to induce hyperkeratosis in the early stages of lymphedema, and this is accompanied by proliferation, inhibited differentiation, and increased expression of stress markers, such as KRT6 (**Fig. 6**).

**Figure 6.**
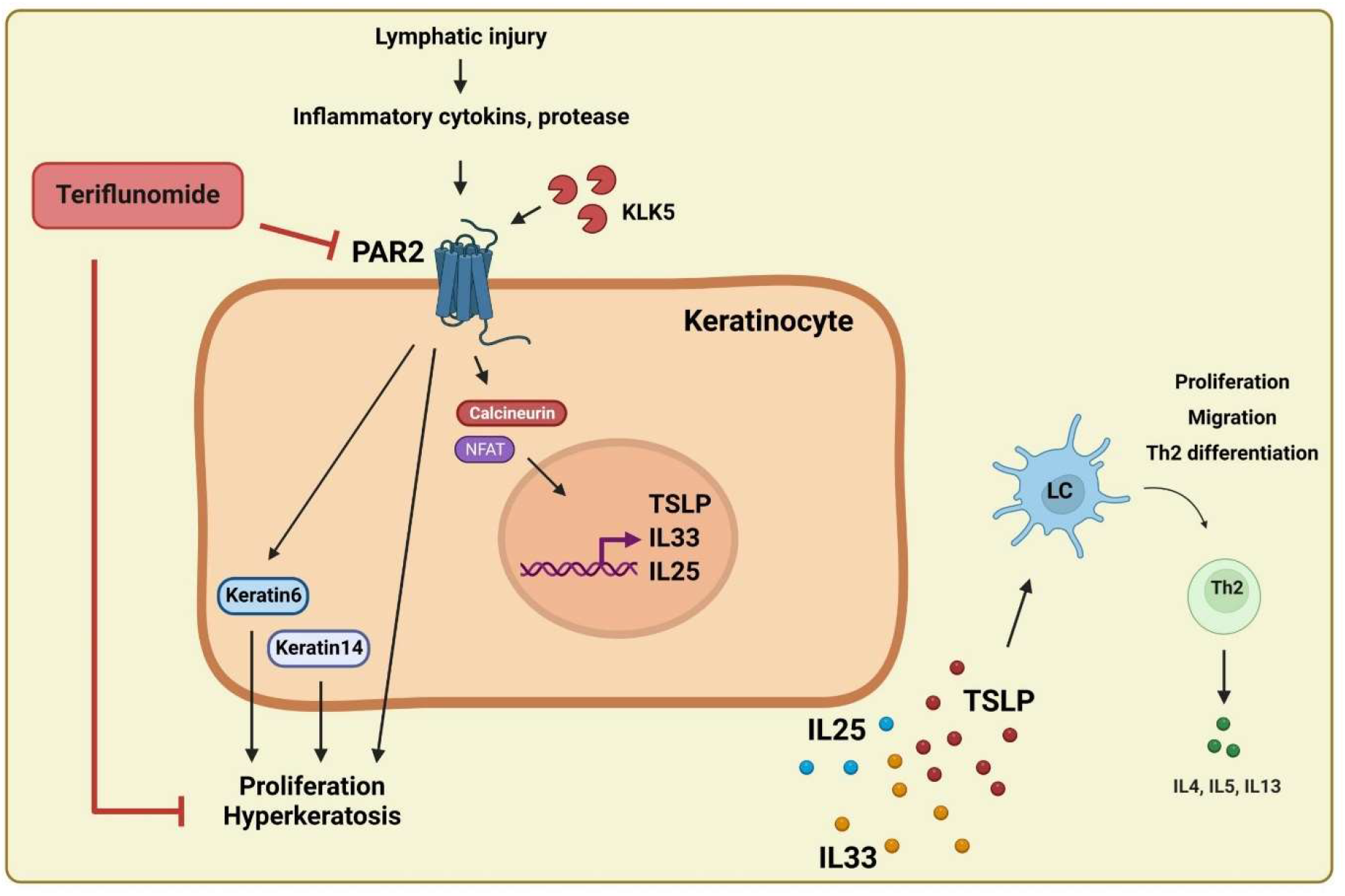
Model of keratinocyte regulated epidermal changes in lymphedema. In the early stages of secondary lymphedema, lymphatic injury induces lymph fluid accumulation and activates hyperkeratosis accompanied by proliferation and increased keratin6 and14 expression. The expression of KLK5-PAR2 and Th2-inducing cytokines (TSLP, IL33, IL25) are upregulated in keratinocytes of lymphedematous skin, which results in Langerhans cell (LC) activation and Th2 differentiation. Topical application of teriflunomide inhibits hyperkeratosis, as well as PAR2-induced Th2-inducing cytokine expression, eventually reducing pathophysiology of secondary lymphedema.

Lymph is fluid that accumulates in interstitial space and is transported by lymphatic vessels. The composition of lymphatic fluid is regulated by physiologic factors, as well as by inflammatory responses elicited by trauma, hemorrhagic shock, or lymphatic injury.(*49*) Tissue injury causes an imbalance in the ratio of protease inhibitor/protease in lymph, resulting in a net activation of protease activity and induction of inflammatory responses.(*50*) Consistent with these observations, we found that the expression of PAR2, a protease receptor, and KLK5, an endogenous skin protease, were significantly increased following lymphatic injury and in lymphedema skin biopsies from patients with unilateral BCRL. PAR2 is a transmembrane protein that is expressed primarily in the skin but also in other tissues, and is activated by proteases including KLK5, trypsin, and papain.(*51*) KLK5 is the most important skin protease that promotes barrier dysfunction in inflammatory skin diseases.(*28, 42*)

We found that PAR2 inhibition with ENMD1068, a small molecule inhibitor, decreased activation of keratinocytes by LF *in vitro* and decreased expression of Th2-inducing cytokines, such as TSLP and IL33. More importantly, inhibition of PAR2 activation in transgenic mice significantly decreased the pathological findings of lymphedema (i.e., fibrosis, swelling) and decreased Th2 inflammatory responses. Loss of PAR2 signaling after skin/lymphatic excision decreased the number of activated LC in regional lymph nodes and the number of infiltrating Th2 cells in the tail skin and draining lymph nodes. Our findings are consistent with previous studies demonstrating that overexpression of KLK5 results in the development of Netherton Syndrome, inflammatory skin lesions, and increased Th2 cell infiltration.(*52*) In contrast, KLK5 inhibition reverses the pathology of Netherton Syndrome.(*28*) Similarly, PAR2 overexpression in basal keratinocytes increases allergic responses against house dust mite, and skin inflammation in mouse models of AD.(*53*) In contrast, PAR2 inactivation reduces early production of TSLP, decreases inflammation and ichthyosis, and decreases Th2 inflammation in mouse models of AD and Netherton Syndrome.(*42, 54, 55*)

We found that, similar to AD and other inflammatory skin disorders, keratinocytes in human and mouse lymphedema samples express Th2-inducing cytokines (TSLP, IL33, and IL25). These findings are important because TSLP, IL33, and IL25 are epithelium-derived cytokines that regulate Th2 inflammation in a variety of settings, including AD, allergic rhinitis, psoriasis, food allergy, and allergic asthma. (*26, 56-58*) Overexpression of TSLP in mouse skin leads to spontaneous AD and increased Th2 cell responses.(*59*) Topical treatment with vitamin D3 analogue MC903 induces AD-like phenotype and TSLP overexpression in keratinocytes, and increased allergic reaction in an allergic asthma mouse model, while KRT14 specific TSLP mutation reversed the effect of MC903.(*57*) Deficiency of TSLP receptor in CD11c^+^ DCs reduces Th2 cell response in a mouse model of lung inflammation, suggesting DCs respond to epithelial-derived TSLP.(*60*) Indeed, TSLP stimulates DC activation by inducing costimulatory molecules, such as OX40L, CD80, and CD86, and TSLP-stimulated DCs promote Th2 polarization.(*26*) Innate immune cells (ILCs) and granulocytes (eosinophil, basophil, mast cells) respond to TSLP and produce inflammatory Th2 cytokines, such as IL4 and IL13.(*26*) IL33 and IL25 also induce Th2 immune responses by acting directly on CD4^+^ T cells, Th2 memory cells, and ILC2s.(*27, 30*) Taken together, substantial evidence suggests that epithelial-derived Th2-inducing cytokines regulate Th2 differentiation and inflammatory disorders and that these responses may also drive inflammatory responses in lymphedema.

The interaction between keratinocytes and Th2 inflammatory cytokines is bidirectional. While keratinocyte-derived Th2-inducing cytokines drive Th2 differentiation, Th2 cytokines, such as IL4 and IL13, in turn regulate keratinocyte differentiation and barrier function.(*61*) *In vitro* treatment of skin equivalent models with Th2 cytokines results in disturbed keratinocyte differentiation and AD-like skin phenotype.(*62*) Th2 cytokines also inhibit expression of skin barrier proteins, such as filaggrin, loricrin, and involucrin, leading to increased skin permeability and sensitivity to bacterial toxins.(*63, 64*) Th2 cytokines also regulate keratinocyte expression of Th2-inducing cytokines, thus acting in a feed-forward manner.(*65*) Our recent clinical trial testing the safety of monoclonal anti-IL4/IL13 antibody treatments in patients with unilateral BCRL are consistent with this paradigm–we observed decreased expression of keratinocyte-derived Th2-inducing cytokines and immune cell recruitment after a 4-month treatment course.(*66*) Changes in skin barrier function related to Th2 cytokine expression may also serve as a putative mechanism for the increased risk of skin infections in some patients with lymphedema and recent reports demonstrating clinical evidence of skin barrier dysfunction and increased transepidermal water loss.(*67*)

Our finding of abnormal keratinocyte proliferation in lymphedema led us to hypothesize that treatment with TF, a proliferation inhibitor, would be an effective treatment for this disease.(*68, 69*) Indeed, we found that *in vitro* treatment of LF-stimulated keratinocytes with TF markedly decreased cellular proliferation and expression of PAR2 and Th2-inducing cytokines. These findings are consistent with previous reports demonstrating that inhibition of PAR2 decreases keratinocyte proliferation, suggesting that PAR2 expression activates a positive feedback response with cellular proliferation.(*70*) It is possible that topical TF used in our mouse models also directly decreases inflammatory responses, since TF can also inhibit proliferation of inflammatory cells. However, the finding that treatment with TF shortly after lymphatic injury (i.e., prior to infiltration of inflammatory cells) also decreased hyperkeratosis and expression of PAR2 suggest that the beneficial effects may be due primarily to the effect of TF on keratinocytes.

Our study has some limitations. Although our mouse lymphedema models closely correlate with the histological and inflammatory changes in BCRL, it is possible that these models do not completely reflect the clinical scenario. However, it is important to note that there are currently no reproducible large animal models of lymphedema—canine models require long-term (>6 month) follow-up after surgery and only result in lymphedema development in a subset of animals;(*71*) pig and sheep models are more consistent with simple lymphatic injury since the procedures do not cause significant, sustained swelling or fibrosis.(*71*) We attempted to mitigate the limitation of mouse models by using a surgical and non-surgical model of lymphedema. Even though the conventional PAR2KO mice used in our study are widely used to study the effects of epidermal PAR2 expression, it is possible that global PAR2 knockout may have effects on inflammatory cells. Thus, future studies will address this issue using tissue-specific knockouts.

In conclusion, our data indicate keratinocytes may play an active role in the development of lymphedema by coordinating Th2 inflammatory responses. Abrogation of this response is highly effective in preclinical models and may represent a novel approach for preventing or treating lymphedema.

## Acknowledgements

Graphical figure is created with BioRender. This research was supported in part by the NIH through R01 HL111130 awarded to B.J.M., T32CA009501 (stipend for G. D. G. N.), and the Cancer Center Support Grant P30 CA008748.

## Author contributions

HJP, RPK and BJM conceived the concept and designed research studies. HJP, RPK and BJM conducted experiments. HJP, JS and GDG acquired data. HJP, GDG, ER and BJM analyzed data. HJP, JS, EME, MGK, MC, JHD and BJM provided reagents and clinical skin biopsies. HJP, RPK and BJM wrote the manuscript.

## Materials and Methods

### Patient samples

All procedures were approved by the Institutional Review Board (IRB protocol 17-377) at Memorial Sloan Kettering Cancer Center (MSK). Women with unilateral upper extremity BCRL were identified in our lymphedema clinic and screened for eligibility for harvesting of biopsy specimens. Inclusion criteria included age between 21-75 years, unilateral axillary surgery, and stage I-III lymphedema (volume differential of >10% with the normal limb or L-Dex measurements above 7.5 units). Exclusion criteria included pregnancy or lactating women, recent (within 3 months) history of lymphedematous limb infection, chemotherapy, treatment with steroids or other immunosuppressive agents, and active cancer or breast cancer metastasis. We harvested excessive interstitial lymph fluid from the lymphedematous arm and mm full-thickness skin biopsies from the volar surface of the normal and lymphedematous arms at a point located 5-10 cm below the elbow crease. Biopsy was performed under sterile conditions with local anesthesia. Patients were treated with a dose of antibiotics (cephalexin 1000 mg or clindamycin 600 mg if penicillin-allergic) 30-60 minutes before the procedure. We obtained informed consent from all patients.

### Animals

All studies were approved by the Institutional Animal Care and Use Committee (IACUC) at MSK (protocol 06-08-018). The MSK IACUC adheres to the National Institutes of Health Public Health Service Policy on Humane Care and Use of Laboratory Animals and operates in accordance with the Animal Welfare Act and the Health Research Extension Act of 1985. Per the IACUC-approved protocol, all mice were maintained in light- and temperature-controlled pathogen-free environments and fed ad libitum.

Adult (8- to 12-week-old) female C57BL/6J mice were used for all treatment studies. We chose to use female mice for our study since secondary lymphedema affects females more commonly than males.(*72*) PAR2 knockout mice (PARKO) based on a C57BL/6 background were purchased from the Jackson Laboratory (B6.Cg-F2rl1^tm1Mslb^/J mice; The Jackson Laboratory, Bar Harbor, ME). Controls were age- and sex-matched wild-type C57B6 mice also purchased from The Jackson Laboratory.

### Surgical model of lymphedema

Anesthesia was induced using isoflurane (Henry Schein Animal Health, Dublin, OH) and mice were kept on a heating blanket to maintain body temperature. Depth of anesthesia was monitored by reaction to pain and observation of respiratory rate. Animals were excluded from the experiment if wound infection or ulceration in the tail was noted at any time point following surgery. Postoperative pain control was maintained with 3 doses of intraperitoneal buprenorphine injection every 4-12 hours. Animals were euthanized by carbon dioxide asphyxiation as recommended by the American Veterinary Medical Association.

In the tail surgery model, both the superficial and deep lymphatic vasculature were ligated through a 2-mm circumferential excision of the skin 2 cm distal to the base of the tail. Collecting lymphatics were identified using Evans Blue injection and ligated along the entire length of the skin excision. Control animals underwent skin incision without lymphatic ligation.(*15, 73*)

### Non-surgical model of lymphedema

Lymphatic-specific diphtheria receptor (DTR) mice were developed as previously described by FLT4-CreERT2 mice (a gift from Sagrario Ortega, CNIO)(*74, 75*) and DTR floxed C57BL/6J mice (C57BL/6-Gt(ROSA)26Sor^tm1(HBEGF)Awai^/J; The Jackson Laboratory). Cre expression was induced in adult mice by using tamoxifen (300 mg/kg/d intraperitoneally for 3 days) followed by local DT (5 ng subcutaneously daily, 3 doses).(*75*) For controls, wild-type C57BL/6J mice were given 3 doses of local subcutaneous DT injection.

### Tail volume measurement

Tail volumes (V) were calculated weekly following tail surgery to evaluate the development of lymphedema over time.(*76*) Digital calipers were used to measure tail diameter every 1 cm starting at the surgical site going distally toward the tip of the tail. Serial circumferences (C) were determined and used to calculate tail volume per the truncated cone formula [V = 1/4π (C_1_C_2_ + C_2_C_3_ + C_3_C_4_)].

### Histology and immunofluorescence

Histological and immunofluorescence analyses were performed using our previously published techniques.(*40*) Clinical and experimental biopsy specimens were fixed in 4% paraformaldehyde (Sigma-Aldrich, St. Louis, MO) overnight. Tails were decalcified using 5% ethylenediaminetetraacetic acid (EDTA; Santa Cruz, Santa Cruz, CA), embedded in paraffin, and sectioned at 5 μm. Hematoxylin and eosin (H&E) staining was performed using standard techniques. For immunofluorescent staining, the rehydrated sections underwent heat-mediated antigen unmasking with sodium citrate (Sigma-Aldrich) and quenching of endogenous peroxidase activity. The sections were then incubated at 4°C with the appropriate primary antibodies overnight. The list of antibodies utilized is in **Table 1**.

**Table 1.** Primary antibodies.

| Target | Origin | Company | Cat. No. |
| --- | --- | --- | --- |
| KRT6 | Mouse | Abcam, Cambridge, UK | ab18586 |
| IL33 | Goat | R&D system, MN, USA | AF3625 |
| KRT14 | Guineapig | Origene, MD, USA | BP5009 |
| Ki67 | Rat | Invitrogen, MA, USA | 14-5689-82 |
| IL1a | Rabbit | Abcam, Cambridge, UK | ab9614 |
| TSLP | Rabbit | Abcam, Cambridge, UK | ab188766 |
| IL33 | Goat | R&D system, MN, USA | AF3626 |
| PAR2 | Rabbit | Abcam, Cambridge, UK | ab180953 |
| KLK5 | Rat | R&D system, MN, USA | MAB7236 |
| NFATc1 | Rabbit | Thermo, MA, USA | PA5-90432 |
| EGF | Rabbit | Abcam, Cambridge, UK | ab9695 |
| EGFR | Rabbit | Abcam, Cambridge, UK | ab52894 |
| Lyve1 | Rabbit | Abcam, Cambridge, UK | ab14917 |
| CD3e | Hamster | Thermo, MA, USA | 14-0031-82 |
| Type I collagen | Rabbit | Abcam, Cambridge, UK | ab34710 |
| Lyve1 | Goat | R&D system, MN, USA | AF2125 |
| CD4 | Rat | R&D system, MN, USA | MAB554 |

H&E and IHC slides were evaluated with brightfield or fluorescent microscopy and scanned using a Mirax slide scanner (Zeiss). Staining was visualized using Pannoramic Viewer (3DHISTECH Ltd., Budapest, Hungary). Epidermal area was quantified in H&E-stained tail cross-sections by measuring the ratio of dark stained epidermis within the total tissue area using MetaMorph Offline software (Molecular Devices, Sunnyvale, CA) with a minimum of 4 high-powered fields per slide by 2 blinded reviewers. Cell counts were quantified in IHC stained tail cross-sections by counting the cells with positive staining. Protein-expressing area was quantified as a ratio of the area of positively stained dermis within a fixed threshold to total tissue area using MetaMorph Offline software (Molecular Devices) with a minimum of 4 high-powered fields per slide by 2 blinded reviewers.

### RNA sequencing

RNA sequencing (RNAseq) was performed in collaboration with the Integrated Genomics Operation (IGO) Core Facility at MSK. Four pairs of frozen clinical lymphedema/normal skin biopsy specimens were submitted to the IGO. The ribodepletion method was used for RNAseq. mRNA expression was standardized and analyzed by IGO. Standardized expression for each molecule was assessed and data are presented as Z-scores. **Real-time qPCR**

Total RNA was extracted using TRIzol (Invitrogen, Carlsbad, CA) according to the manufacturer’s instruction, and complementary DNA (cDNA) was prepared by using Maxima™ H Minus cDNA Synthesis Master Mix (Thermo Scientific, Rockford, IL). Real-time qPCR (qRT-PCR; ViiA7; Life Technologies, Carlsbad, CA) was performed in duplicates using predesigned primer sets (Quantitect Primer Assays, Qiagen, Germantown, MD). Relative mRNA expression between groups was analyzed by the delta-delta Ct method and normalized to housekeeping genes, β-actin or GAPDH. Standardized expression for each molecule was assessed, and data are presented as Z-scores.

### Western blot

Clinical and mouse skin biopsies were frozen in liquid nitrogen, homogenized, and lysed with a radioimmunoprecipitation assay (RIPA) lysis buffer containing Halt Protease and Phosphatase Inhibitor Cocktail (Thermo Fisher Scientific, Waltham, MA). The lysates were centrifuged at 13,000 x g for 10 minutes at 4°C, and protein concentration was measured by BCA protein assay kit (Thermo Fisher Scientific) according to manufacturer’s directions. One to 20 μg of total protein were separated by NuPAGE™ 4-12% Bis-Tris Gel (Thermo Fisher Scientific) and transferred onto PVDF membranes (Bio-Rad, Hercules, CA). Membranes were blocked with 5% skim milk in TBS containing 0.1% Tween 20 (TBST) at room temperature for 1 hour and incubated with antibodies against KRT6 (ab18586; Abcam), KRT16 (ab154361; Abcam), PAR2 (ab180953; Abcam), KLK5 (MAB7236; R&D system), β-actin (3700s; Cell signaling) in 0.5% skim milk in TBST at 4°C overnight. After washing 3 times with TBST, membranes were incubated with HRP-conjugated secondary antibody in TBST at room temperature for 1 hour. Then, membranes were washed with TBST, and immune-reactive bands were detected with ECL Western Blotting Substrate (Thermo Fisher Scientific). Protein expression was quantified with ImageJ software (National Institutes of Health, Bethesda, MD) and normalized to housekeeping genes, GAPDH or β-actin.

### ELISA

ELISA was performed using our published methods.(*8*) Briefly, tail skin tissue was harvested 1.5 cm distal to the surgical site, flash-frozen in liquid nitrogen, and protein was extracted with tissue extraction protein reagent (ThermoFisher Scientific) mixed with phosphatase and protease inhibitor (Sigma-Aldrich). Approximately 20-30 mg of protein from each sample was analyzed per the manufacturer’s recommendations for each protein. The following ELISA kits were used: TSLP mouse ELISA kit (EMTSLP; Thermo) and Mouse IL-33 Quantikine ELISA kit (M3300; R&D system). All samples were assessed in triplicate.

### Flow cytometry

Flow cytometry was performed to quantify inflammation in the mouse tails after tail surgery.(*16*) Briefly, single-cell suspensions were obtained from a 1-cm portion of the tail distal to the surgical site using a combination of mechanical dissociation and enzymatic digestion with a solution of DNase I, Dispase II, collagenase D, and collagenase IV (all Roche Diagnostics; Indianapolis, IN) mixed in 2% fetal calf serum (FCS; Sigma-Aldrich). Cells were stained with combinations of the following fluorophore-conjugated anti-mouse monoclonal antibodies: Rat CD45 (30-F11; #103139), Rat CD45 (30-F11; #103116), Rat CD11b (M1/70; #101228), Armenian hamster CD11c (N418; #117306), Mouse CD207 (4C7; #144206), Rat CD4 (GK1.5; #100408), Armenian hamster CXCR3 (CXCR3-173; #126536), Armenian hamster CCR5 (HM-CCR5; #107016), Armenian hamster CCR4 (2G12; #131214), Rat CCR8 (SA214G2; #150310) from BioLegened (San Diego, CA); Rat F4/80 (BM8; #25-4801-82) from eBioscience (San Diego, CA). In addition, DAPI viability stain was also used on all samples to exclude dead cells. Single-stain compensation samples were created using UltraComp eBeads™ (#01-2222-42, Affymetrix, Inc.; San Diego, CA). Flow cytometry was performed using a BD Fortessa flow cytometry analyzer (BD Biosciences) with BD FACS Diva, and data were analyzed with FlowJo software (Tree Star; Ashland, OR).

### *In vitro* keratinocyte culture and treatment

Human keratinocytes (PCS-200-011; ATCC, VA) were cultured in dermal cell basal medium (PCS-200-030; ATCC) with keratinocyte growth kit (PCS-200-040; ATCC). Keratinocytes were cultured with or without 10% lymphedema fluid in keratinocyte medium. ENMD1068, a PAR2 antagonist (ab141699; Abcam; 10 µg/ml); was added once to the culture media at the same time as the lymphedema fluid treatment. Teriflunomide (TF; 25 µM in DMSO) or DMSO alone were added to the culture media at the same time as the lymphedema fluid treatment. Cells were harvested 6 or 48 hours after the treatment for RNA or protein extraction, respectively. Proliferation of keratinocytes was measured by using Vybrant MTT cell proliferation assay kit (V13154; Thermo). Teriflunomide (indicated concentration in DMSO) or DMSO were added to the culture media a day after seeding for MTT assay and cultured for 24 hours.

### Topical teriflunomide treatment

A topical formulation of TF(27 mg/ml dissolved in Aquaphor®; Beiersdorf, Hamburg, Germany) was developed in collaboration with the MSK Research Pharmacy Core Facility. Control animals were treated with Aquaphor® alone. The treatment was applied once daily for 2 or 4 weeks to the tail region distal to the zone of lymphatic skin excision or for 8 weeks to the footpad.

### Statistical analysis

Statistical analyses were performed using GraphPad Prism 9.0.2 (GraphPad Software, Inc.; San Diego, CA). Samples were assessed for normal distribution using the Shapiro Wilk test. Normally distributed clinical samples were analyzed using paired student’s t-test. Comparisons of multiple groups or time points were performed using unpaired student’s t-test or Mann-Whitney test or one-way ANOVA or two-way ANOVA with multiple comparisons using Tukey’s multiple comparison test. Data are presented as mean ± standard deviation unless otherwise noted, with *p*<0.05 considered significant. For all plots, each dot represents one animal or patient unless noted otherwise.

---

**Figure S1.**
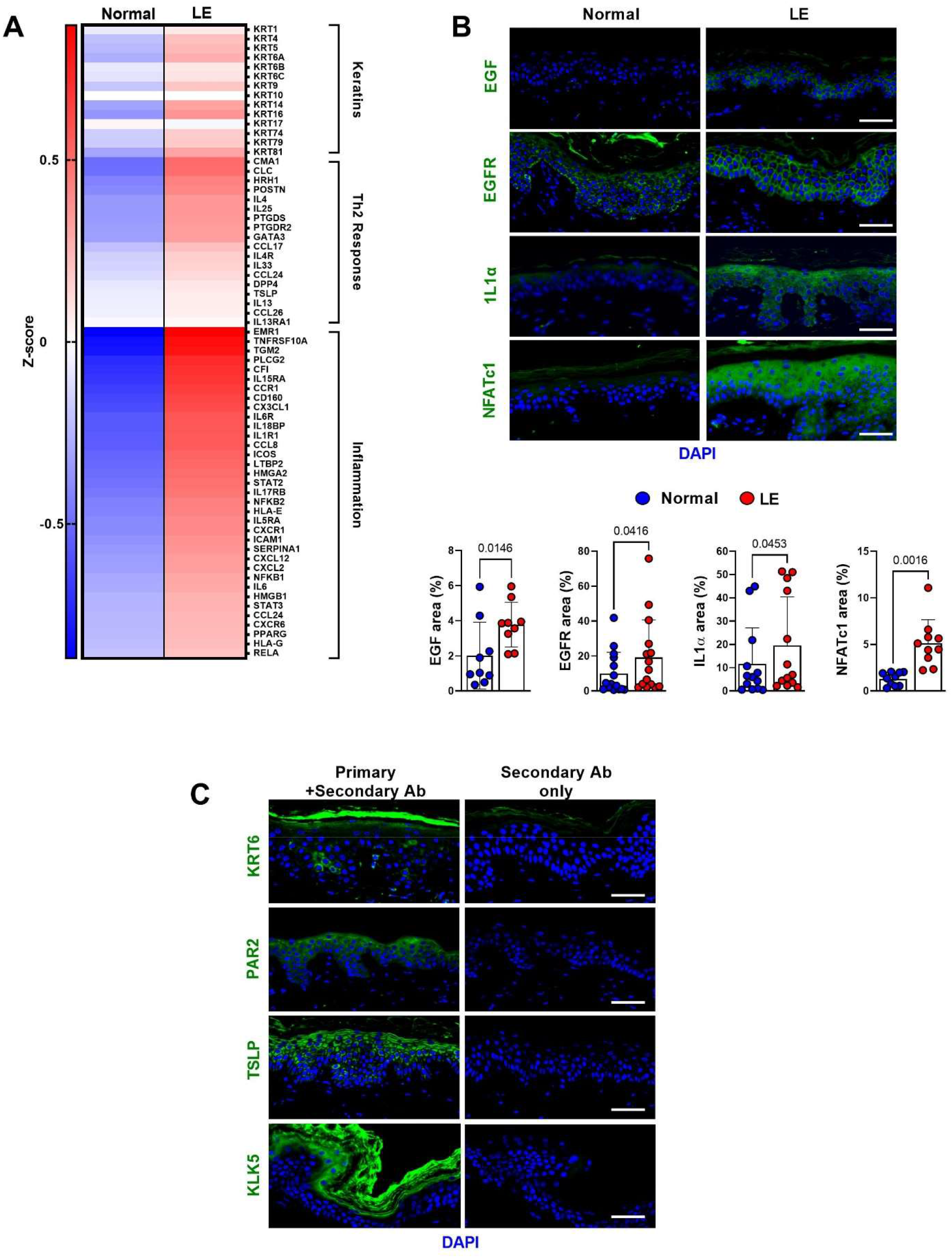
Expression of keratinocyte growth factors and inflammatory cytokines are upregulated in lymphedema. A. Expression of keratins, Th2 response, and inflammatory genes by RNAseq in normal and lymphedematous (LE) skin biopsies from patients with unilateral BCRL (N=4). Each box represents the average mRNA expression of 4 patients. B. Representative immunofluorescent images (*top*) and quantification (*bottom*) of EGF, EGFR, IL1α, and NFATc1 area in normal and lymphedematous (LE) skin biopsies from patients with unilateral BCRL. Scale bar: 50 µm. Each circle represents the average quantification of 3 HPF views for each patient (N=10-15). *P* values were calculated by paired student’s t-test. C. Immunofluorescent analysis of lymphedema skin biopsies with each antibody (Ab) and respective negative controls without primary antibodies. Scale bar: 50 µm.

**Figure S2.**
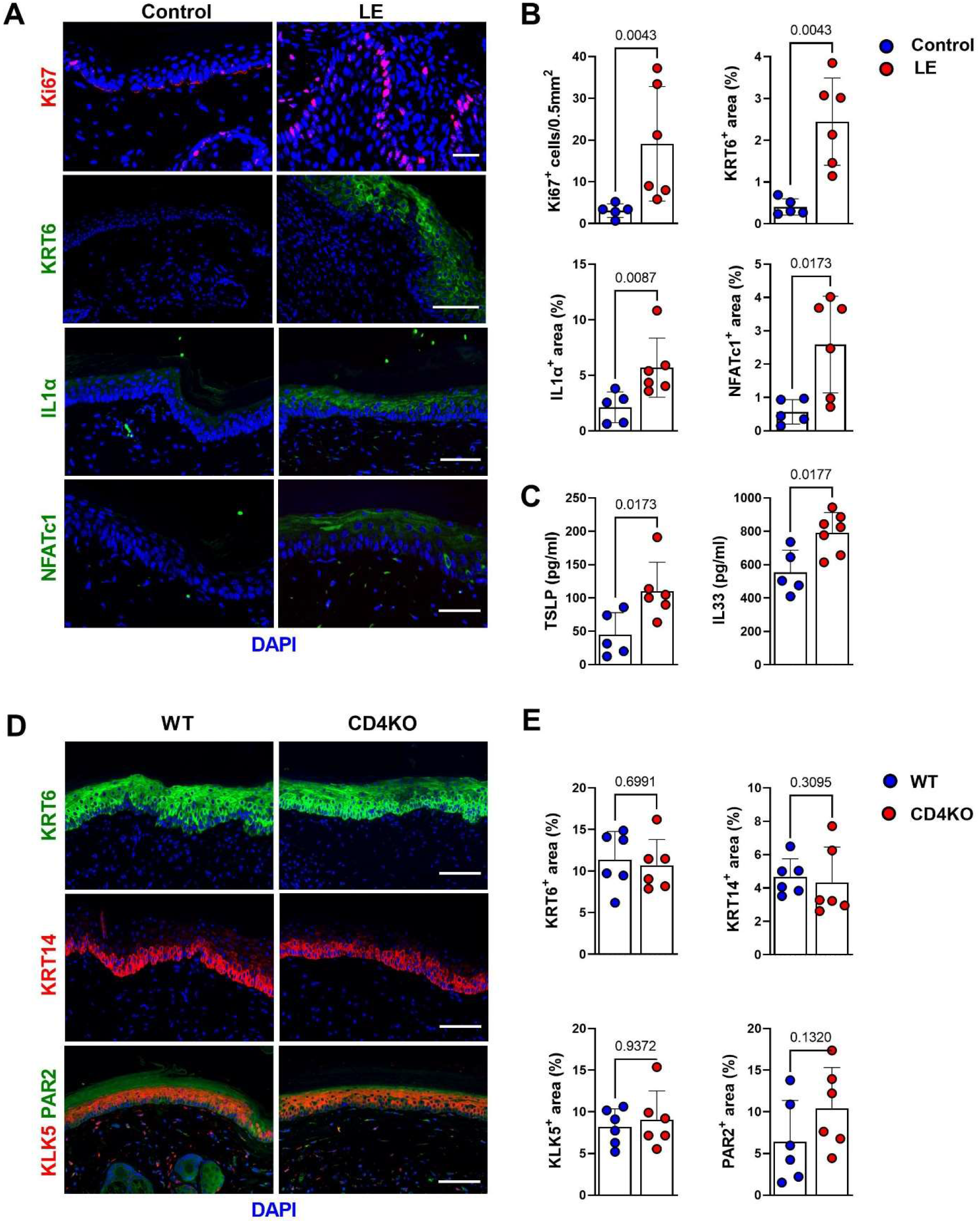
Gene expression changes in keratinocytes occur in rapidly after lymphatic injury and are independent of CD4 T cells. A. Representative immunofluorescent images of KRT6, Ki67, IL1α, and NFATc1 staining in tail skin harvested 2 weeks after surgery from control and lymphedema (LE) mice. Scale bar: 100 µm. B. Quantification of KRT6, Ki67, IL1α, and NFATc1 area in tail skin harvested 2 weeks after surgery from control and lymphedema (LE) mice. Each circle represents the average of 3 HPF views of each mouse (N=5-6). *P* values were calculated by Mann-Whitney test. C. TSLP and IL33 ELISA from protein lysates of tail skin harvested 2 weeks after surgery from control and lymphedema (LE) mice (N=5-7). Each circle represents one mouse. *P* values were calculated by Mann-Whitney test. D. Representative immunofluorescent images of KRT6, KRT14, KLK5, and PAR2 staining in tail specimens harvested 2 weeks after tail skin and lymphatic excision in wild-type (WT) and CD4 knockout (CD4KO) mice. Scale bar: 100 µm. E. Quantification of KRT6, KRT14, KLK5, and PAR2 area in tail specimens harvested 2 weeks after tail skin and lymphatic excision in wild-type (WT) and CD4 knockout (CD4KO) mice. Each circle represents the average of 3 HPF views of each mouse (N=6).

**Figure S3.**
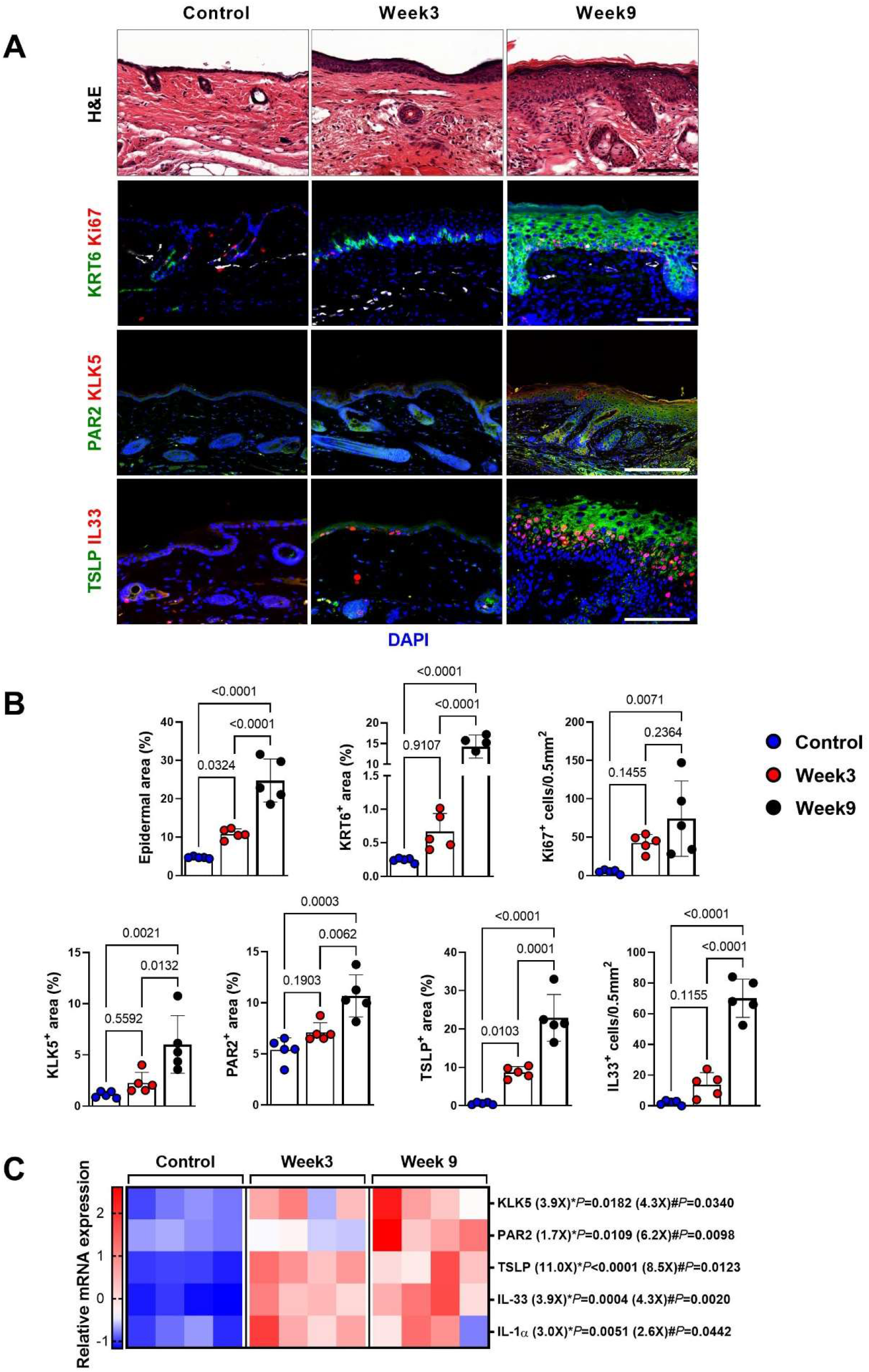
Hyperkeratosis and Th2-inducing cytokine expression are increased in a non-surgical model of lymphedema. A. Representative H&E and immunofluorescent images of KRT6, Ki67, KLK5, PAR2, TSLP, and IL33 in WT hindlimb skin harvested 9 weeks after DT injection and FLT4CreDTRfloxed hindlimb skin harvested 3 or 9 weeks after DT injection. Scale bar: 100 µm. B. Quantification of epidermal, KRT6, Ki67, KLK5, PAR2, TSLP, and IL33 area in WT hindlimb skin harvested 9 weeks after DT injection and FLT4CreDTRfloxed hindlimb skin harvested 3 or 9 weeks after DT injection. Each circle represents the average quantification of 3 HPF views for each mouse (N=5). *P* values were calculated by one-way ANOVA. C. Relative mRNA expression by qPCR in WT hindlimb skin harvested 9 weeks after DT injection and FLT4CreDTRfloxed hindlimb skin harvested 3 or 9 weeks after DT injection. (N=4). mRNA expression was normalized to β-actin expression. Each box represents one mouse. \**P* indicates week 3 compared to control; #*P* indicates week 9 compared to control. *P* values were calculated by Mann-Whitney test.

**Figure S4.**
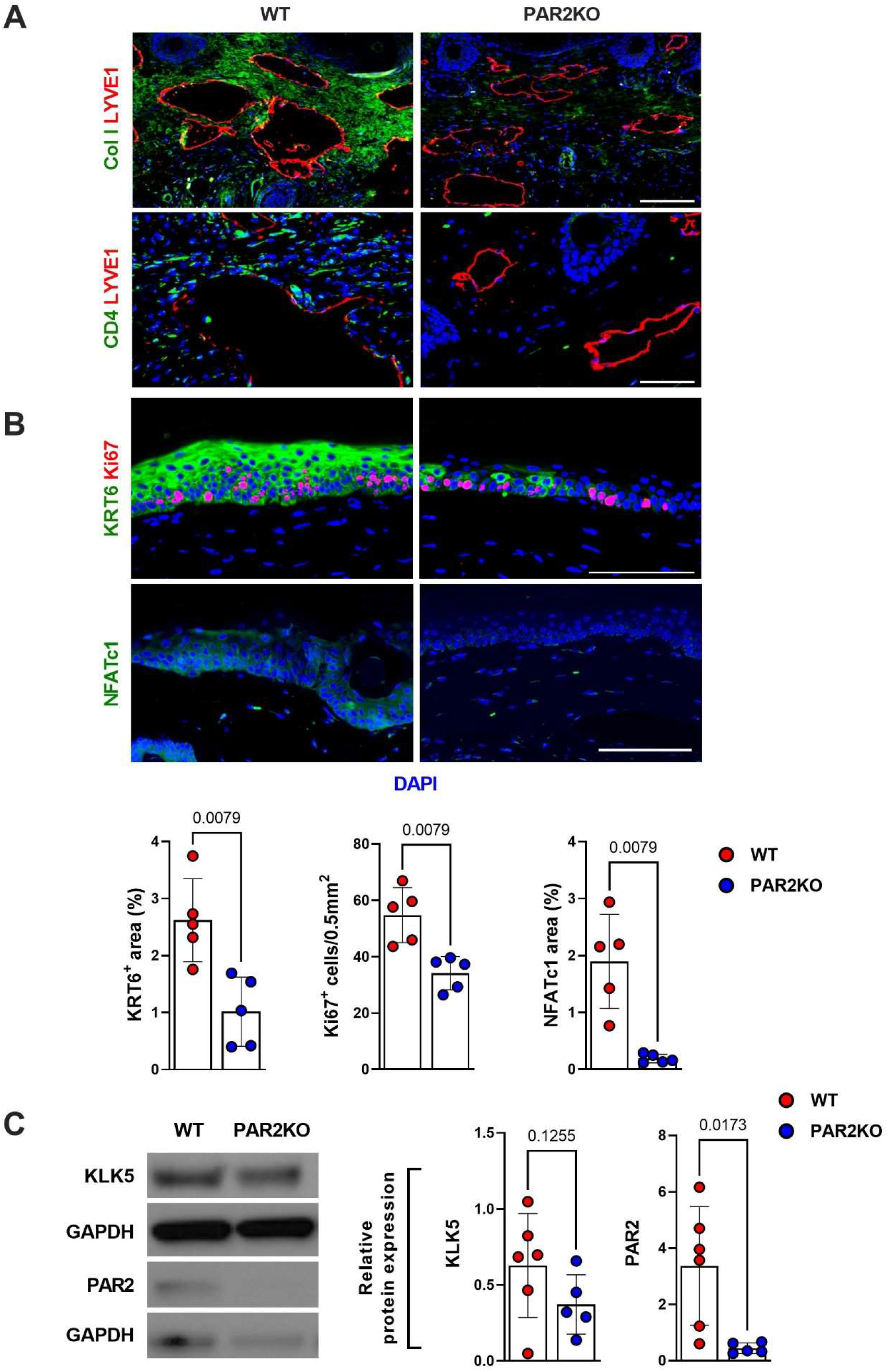
PAR2 knockout decreases fibrosis, CD4^+^ cell infiltration, and hyperkeratosis after lymphatic injury. A. Representative immunofluorescent images of collagen I, LYVE1, and CD4 staining in lymphedema tail skin of wild-type (WT) and PAR2 knockout (PAR2KO) mice harvested 6 weeks after tail skin and lymphatic excision. Scale bar: 100 µm. B. Representative immunofluorescent images (*top*) and quantification (*bottom*) of KRT6, Ki67, and NFATc1 area in tail skin of WT and PAR2KO mice harvested 6 weeks after tail skin and lymphatic excision. Scale bar: 100 µm. Each circle represents the average quantification of 3 HPF views for each mouse (N=5; *bottom*). *P* values were calculated by Mann-Whitney test. C. Representative western blots of KLK5 and PAR2 in lymphedema tail skin of WT and PAR2KO mice harvested 6 weeks after tail skin and lymphatic excision (*left*) and quantification relative to GAPHD (*right*). Each circle represents each mouse (N=5-6). *P* values were calculated by Mann-Whitney test.

**Figure S5.**
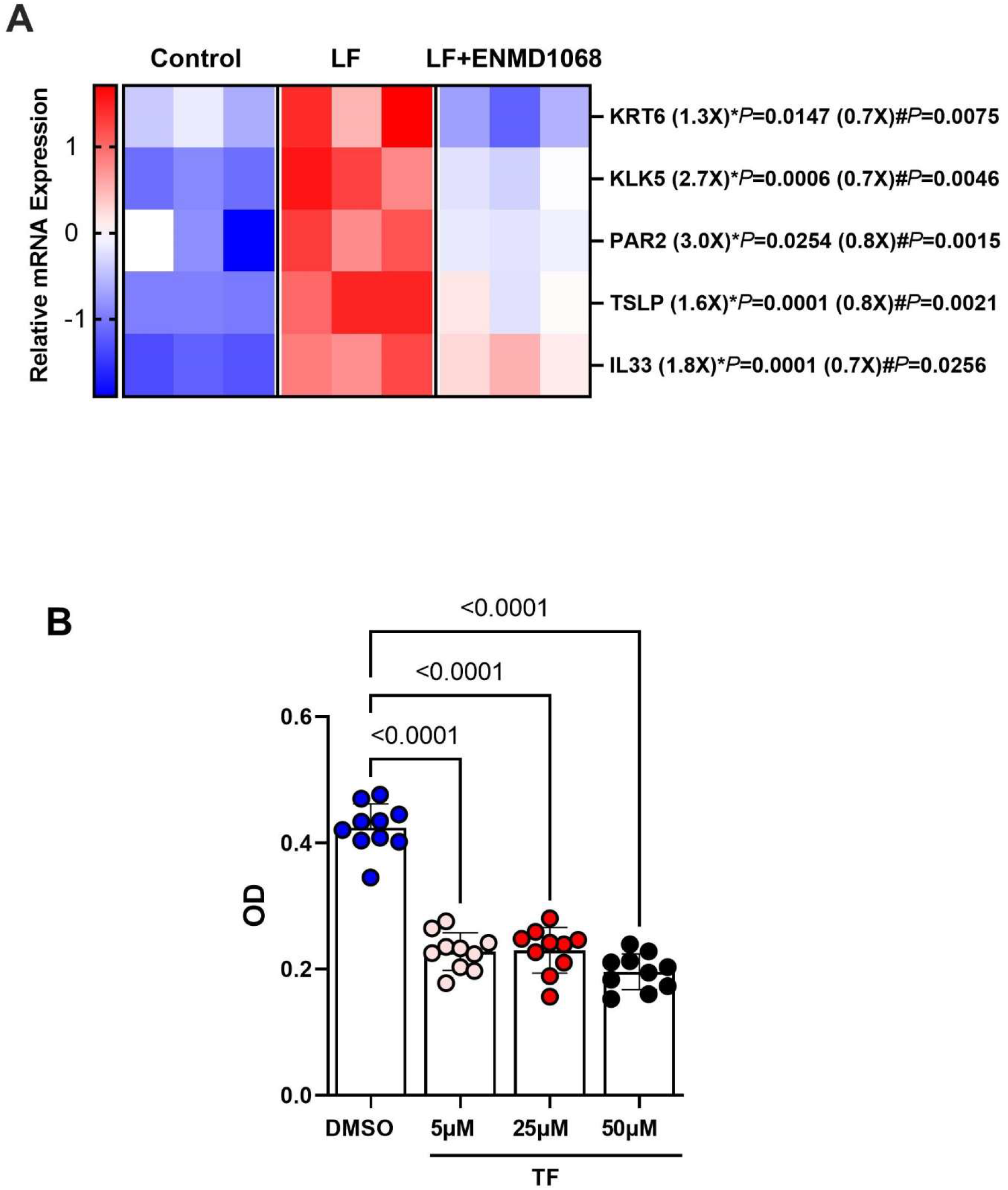
ENMD1068 and teriflunomide decrease keratinocyte activation in response to lymphatic fluid. A. Relative mRNA expression by qPCR in cultured h-keratinocytes treated with PBS (control), lymphatic fluid (LF), or LF+ENMD (N=3). mRNA expression was normalized by β-actin expression. Each box represents each experiment with independently cultured keratinocytes. Fold changes relative to control for LF, and relative to LF for LF+ENMD. \**P* indicates LF compared to control, #*P* indicates for LF+ENMD compared to LF. *P* values were calculated by Mann-Whitney test. B. Proliferation (MTT assay) of h-keratinocytes cultured with DMSO only or teriflunomide (TF) at the indicated concentrations (N=10). Each circle represents an individual experiment. *P* values were calculated by one-way ANOVA.

**Figure S6.**
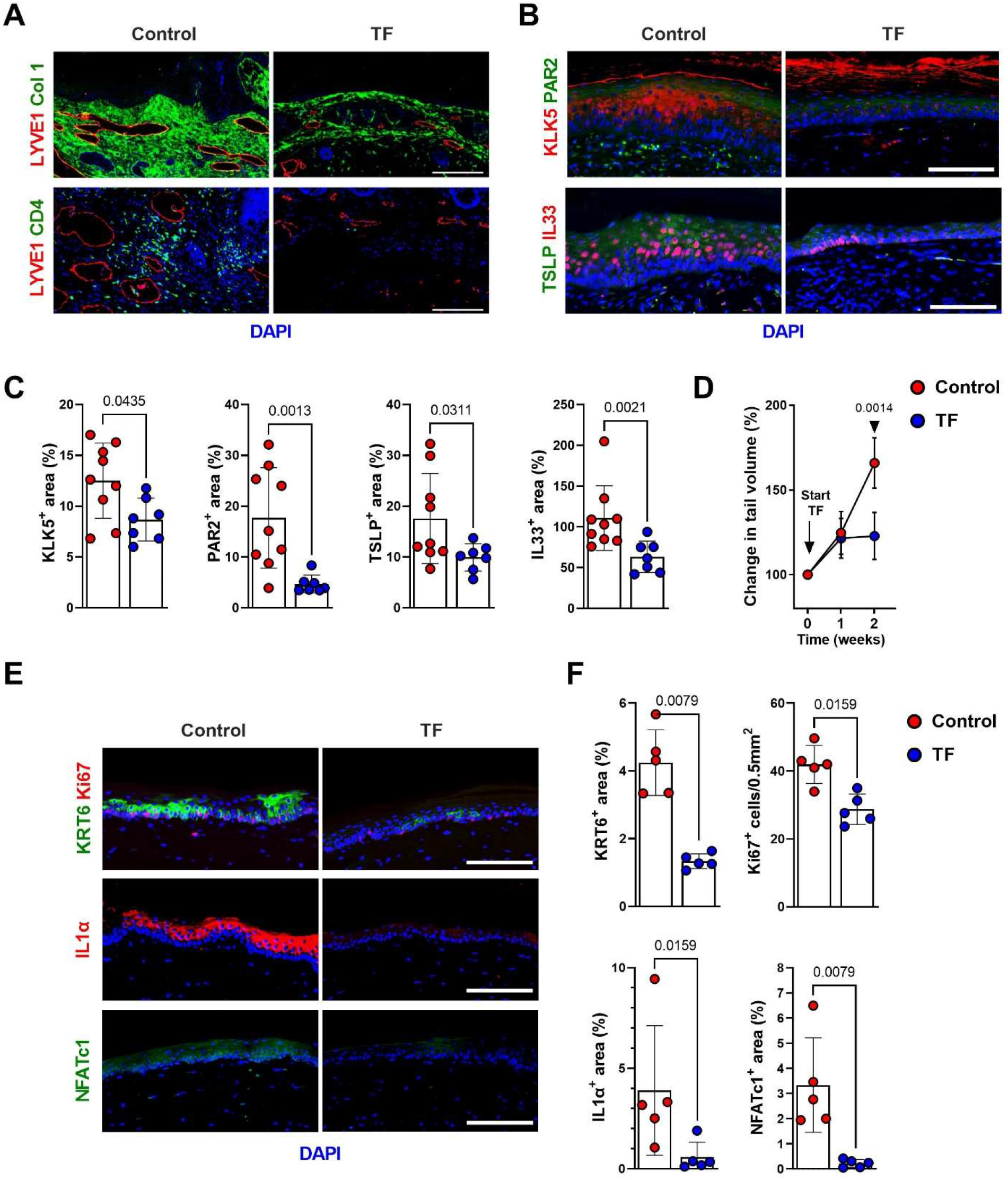
Teriflunomide treatment decreases fibrosis, CD4^+^ infiltration, and hyperkeratosis after lymphatic injury. A. Representative immunofluorescent images of collagen I, LYVE1, and CD4 staining in tail skin samples harvested from mice treated with vehicle (control) or teriflunomide (TF) once daily for 4 weeks beginning 2 weeks after tail skin and lymphatic excision. Scale bar: 100 µm. B. Representative immunofluorescent images of KLK5, PAR2, TSLP, and IL33 staining in tail skin samples harvested from mice treated with vehicle (control) or teriflunomide (TF) once daily for 4 weeks beginning 2 weeks after tail skin and lymphatic excision. Scale bar: 100 µm. C. Quantification of KLK5, PAR2, TSLP, and IL33 area in vehicle and TF-treated mice. Each circle represents the average quantification of 3 HPF views for each mouse (N=7-9). *P* values were calculated by Mann-Whitney test. D. Changes in tail volume over time in mice treated with vehicle (control) or TF for 2 weeks starting one day after tail skin and lymphatic excision. Each circle represents the average measurement from each mouse (N=5). *P* values were calculated by 2-way ANOVA with multiple comparisons. E. Representative immunofluorescent images of KRT6, Ki67, IL1α, and NFATc1 staining. of tail skin from mice treated with vehicle (control) or TF for 2 weeks starting one day after tail skin and lymphatic excision. Scale bar: 100 µm. F. Quantification of KRT6, Ki67, IL1α, and NFATc1 area in vehicle and TF-treated mice. Each circle represents the average quantification of 3 HPF views for each mouse (N=5). *P* values were calculated by Mann-Whitney test.

**Figure S7.**
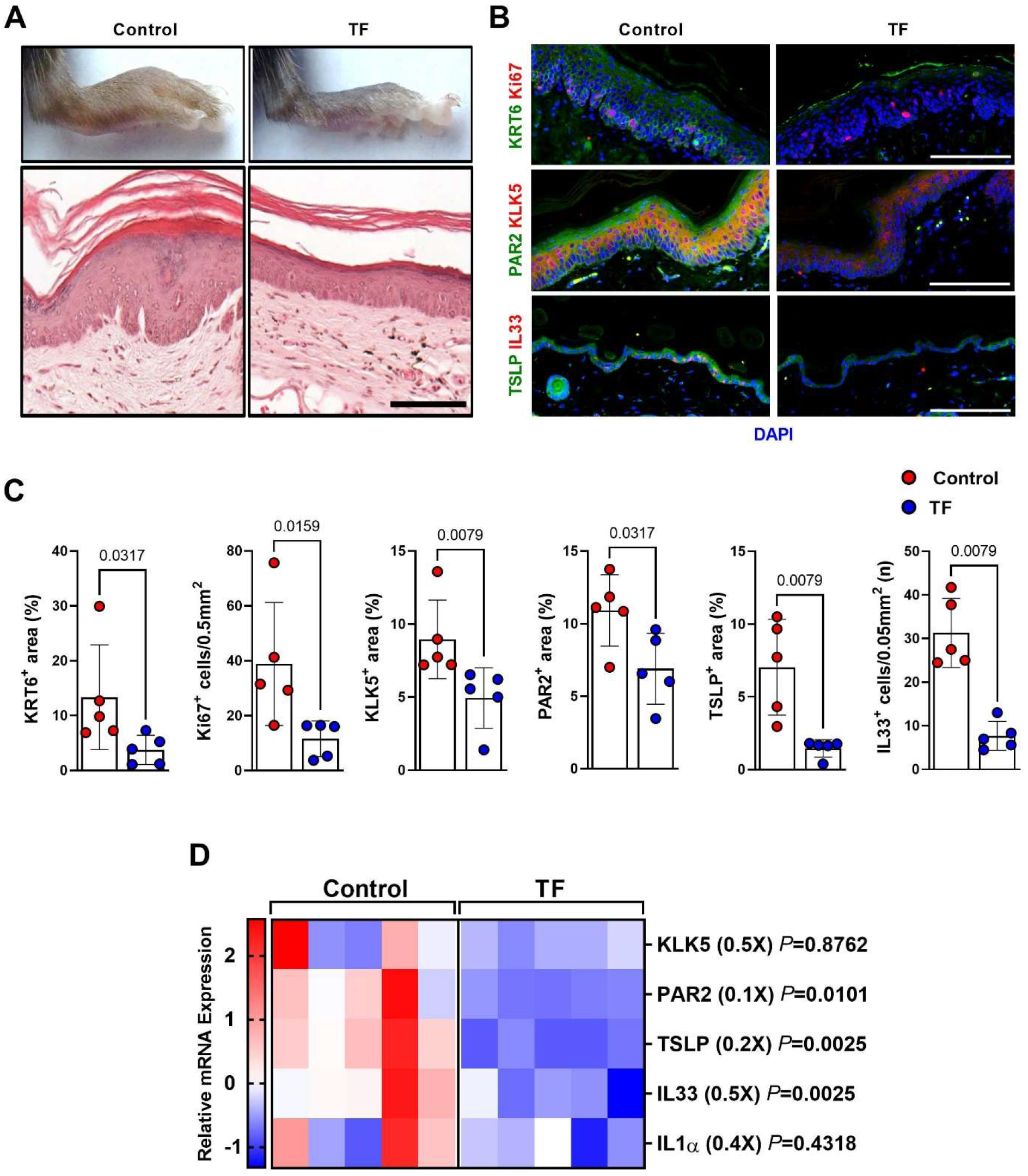
Teriflunomide decreases lymphedema in a non-surgical mouse model. A. Representative images of lymphedema hindlimb and H&E images of lymphedema hindlimb skin from mice treated with vehicle (control) or teriflunomide (TF) for 8 weeks starting a week after DT injection. Scale bar: 100 µm. B. Representative immunofluorescent images of KRT6, Ki67, KLK5, PAR2, TSLP, and IL33 staining of hindlimb skin from control and TF-treated mice. Scale bar: 100 µm. C. Quantification of KRT6, Ki67, KLK5, PAR2, TSLP, and IL33 area in hindlimb skin from control and TF-treated mice. Each circle represents the average quantification of 3 HPF views for each mouse (N=5). *P* values were calculated by Mann-Whitney test. D. Relative mRNA expression by qPCR in hindlimb skin from vehicle (control) and TF-treated mice (N=5). mRNA expression was normalized to β-actin expression. Each box represents one mouse. *P* values were calculated by Mann-Whitney test. Fold change from control is shown in parentheses.

## References

1. A. Szuba, S. G. Rockson, Lymphedema: anatomy, physiology and pathogenesis. Vasc Med 2, 321–326 (1997).

2. J. A. Petrek, M. C. Heelan, Incidence of breast carcinoma-related lymphedema. Cancer 83, 2776–2781 (1998).

3. J. H. Dayan, C. L. Ly, R. P. Kataru, B. J. Mehrara, Lymphedema: Pathogenesis and Novel Therapies. Annu Rev Med 69, 263–276 (2018).

4. S. Vignes, R. Porcher, M. Arrault, A. Dupuy, Long-term management of breast cancer-related lymphedema after intensive decongestive physiotherapy. Breast Cancer Res Treat 101, 285–290 (2007).

5. D. W. Chang et al., Surgical Treatment of Lymphedema: A Systematic Review and Meta-Analysis of Controlled Trials. Results of a Consensus Conference. Plast Reconstr Surg 147, 975–993 (2021).

6. C. L. Ly, D. A. Cuzzone, R. P. Kataru, B. J. Mehrara, Small Numbers of CD4+ T Cells Can Induce Development of Lymphedema. Plast Reconstr Surg 143, 518e–526e (2019).

7. G. D. García Nores et al., CD4(+) T cells are activated in regional lymph nodes and migrate to skin to initiate lymphedema. Nat Commun 9, 1970 (2018).

8. J. C. Gardenier et al., Topical tacrolimus for the treatment of secondary lymphedema. Nat Commun 8, 14345 (2017).

9. F. Ogata et al., Excess Lymphangiogenesis Cooperatively Induced by Macrophages and CD4(+) T Cells Drives the Pathogenesis of Lymphedema. J Invest Dermatol 136, 706–714 (2016).

10. E. Gousopoulos et al., Regulatory T cell transfer ameliorates lymphedema and promotes lymphatic vessel function. JCI Insight 1, e89081 (2016).

11. Y. Yuan, V. Arcucci, S. M. Levy, M. G. Achen, Modulation of Immunity by Lymphatic Dysfunction in Lymphedema. Front Immunol 10, 76 (2019).

12. H. Hara et al., Pathological Investigation of Acquired Lymphangiectasia Accompanied by Lower Limb Lymphedema: Lymphocyte Infiltration in the Dermis and Epidermis. Lymphat Res Biol 14, 172–180 (2016).

13. K. Nakamura, K. Radhakrishnan, Y. M. Wong, S. G. Rockson, Anti-inflammatory pharmacotherapy with ketoprofen ameliorates experimental lymphatic vascular insufficiency in mice. PLoS One 4, e8380 (2009).

14. S. G. Rockson et al., Pilot studies demonstrate the potential benefits of antiinflammatory therapy in human lymphedema. JCI Insight 3, (2018).

15. T. Avraham et al., Th2 differentiation is necessary for soft tissue fibrosis and lymphatic dysfunction resulting from lymphedema. FASEB J 27, 1114–1126 (2013).

16. J. C. Zampell et al., CD4(+) cells regulate fibrosis and lymphangiogenesis in response to lymphatic fluid stasis. PLoS One 7, e49940 (2012).

17. S. Ghanta et al., Regulation of inflammation and fibrosis by macrophages in lymphedema. American journal of physiology. Heart and circulatory physiology 308, H1065–1077 (2015).

18. C. L. Ly, G. D. G. Nores, R. P. Kataru, B. J. Mehrara, T helper 2 differentiation is necessary for development of lymphedema. Transl Res 206, 57–70 (2019).

19. I. L. Savetsky et al., Th2 cytokines inhibit lymphangiogenesis. PLoS One 10, e0126908 (2015).

20. B. Mehrara et al., Pilot study of anti-IL4/IL13 immunotherapy for the treatment of breast cancer–related upper extremity lymphedema. Submitted for publication (Cancers) In press, (2021).

21. J. Furlong-Silva et al., Tetracyclines improve experimental lymphatic filariasis pathology by disrupting interleukin-4 receptor-mediated lymphangiogenesis. J Clin Invest 131, (2021).

22. J. Horton et al., The design and development of a multicentric protocol to investigate the impact of adjunctive doxycycline on the management of peripheral lymphoedema caused by lymphatic filariasis and podoconiosis. Parasit Vectors 13, 155 (2020).

23. A. Domaszewska-Szostek, M. Zaleska, W. L. Olszewski, Hyperkeratosis in human lower limb lymphedema: the effect of stagnant tissue fluid/lymph. J Eur Acad Dermatol Venereol 30, 1002–1008 (2016).

24. H. E. De Cock, V. K. Affolter, E. R. Wisner, G. L. Ferraro, N. J. MacLachlan, Progressive swelling, hyperkeratosis, and fibrosis of distal limbs in Clydesdales, Shires, and Belgian draft horses, suggestive of primary lymphedema. Lymphat Res Biol 1, 191–199 (2003).

25. M. Carretero et al., Differential Features between Chronic Skin Inflammatory Diseases Revealed in Skin-Humanized Psoriasis and Atopic Dermatitis Mouse Models. J Invest Dermatol 136, 136–145 (2016).

26. F. Roan, K. Obata-Ninomiya, S. F. Ziegler, Epithelial cell-derived cytokines: more than just signaling the alarm. J Clin Invest 129, 1441–1451 (2019).

27. R. Divekar, H. Kita, Recent advances in epithelium-derived cytokines (IL-33, IL-25, and thymic stromal lymphopoietin) and allergic inflammation. Curr Opin Allergy Clin Immunol 15, 98–103 (2015).

28. L. Furio et al., Transgenic kallikrein 5 mice reproduce major cutaneous and systemic hallmarks of Netherton syndrome. J Exp Med 211, 499–513 (2014).

29. V. Soumelis et al., Human epithelial cells trigger dendritic cell mediated allergic inflammation by producing TSLP. Nat Immunol 3, 673–680 (2002).

30. Y. H. Wang et al., IL-25 augments type 2 immune responses by enhancing the expansion and functions of TSLP-DC-activated Th2 memory cells. J Exp Med 204, 1837–1847 (2007).

31. M. Omori, S. Ziegler, Induction of IL-4 expression in CD4(+) T cells by thymic stromal lymphopoietin. J Immunol 178, 1396–1404 (2007).

32. H. F. Rosenberg, K. D. Dyer, P. S. Foster, Eosinophils: changing perspectives in health and disease. Nat Rev Immunol 13, 9–22 (2013).

33. J. J. McLeod, B. Baker, J. J. Ryan, Mast cell production and response to IL-4 and IL-13. Cytokine 75, 57–61 (2015).

34. A. Chiricozzi, M. Maurelli, K. Peris, G. Girolomoni, Targeting IL-4 for the Treatment of Atopic Dermatitis. Immunotargets Ther 9, 151–156 (2020).

35. A. S. Rothmeier, W. Ruf, Protease-activated receptor 2 signaling in inflammation. Semin Immunopathol 34, 133–149 (2012).

36. S. S. Athari, Targeting cell signaling in allergic asthma. Signal Transduct Target Ther 4, 45 (2019).

37. X. Zhang, M. Yin, L. J. Zhang, Keratin 6, 16 and 17-Critical Barrier Alarmin Molecules in Skin Wounds and Psoriasis. Cells 8, (2019).

38. H. Alam, L. Sehgal, S. T. Kundu, S. N. Dalal, M. M. Vaidya, Novel function of keratins 5 and 14 in proliferation and differentiation of stratified epithelial cells. Mol Biol Cell 22, 4068–4078 (2011).

39. A. Jairaman, M. Yamashita, R. P. Schleimer, M. Prakriya, Store-Operated Ca2+ Release-Activated Ca2+ Channels Regulate PAR2-Activated Ca2+ Signaling and Cytokine Production in Airway Epithelial Cells. J Immunol 195, 2122–2133 (2015).

40. J. E. Baik et al., TGF-beta1 mediates pathologic changes of secondary lymphedema by promoting fibrosis and inflammation. Clin Transl Med 12, e758 (2022).

41. J. C. Gardenier et al., Diphtheria toxin-mediated ablation of lymphatic endothelial cells results in progressive lymphedema. JCI Insight 1, e84095 (2016).

42. S. Frateschi et al., PAR2 absence completely rescues inflammation and ichthyosis caused by altered CAP1/Prss8 expression in mouse skin. Nat Commun 2, 161 (2011).

43. L. Furio et al., KLK5 Inactivation Reverses Cutaneous Hallmarks of Netherton Syndrome. PLoS Genet 11, e1005389 (2015).

44. M. D. Wiese, A. Rowland, T. M. Polasek, M. J. Sorich, C. O’Doherty, Pharmacokinetic evaluation of teriflunomide for the treatment of multiple sclerosis. Expert Opin Drug Metab Toxicol 9, 1025–1035 (2013).

45. K. Ruckemann et al., Leflunomide inhibits pyrimidine de novo synthesis in mitogen-stimulated T-lymphocytes from healthy humans. J Biol Chem 273, 21682–21691 (1998).

46. T. Matsui, M. Amagai, Dissecting the formation, structure and barrier function of the stratum corneum. Int Immunol 27, 269–280 (2015).

47. R. Gniadecki, Regulation of keratinocyte proliferation. Gen Pharmacol 30, 619–622 (1998).

48. S. Werner, H. Smola, Paracrine regulation of keratinocyte proliferation and differentiation. Trends Cell Biol 11, 143–146 (2001).

49. E. Oveland et al., Proteomic evaluation of inflammatory proteins in rat spleen interstitial fluid and lymph during LPS-induced systemic inflammation reveals increased levels of ADAMST1. J Proteome Res 11, 5338–5349 (2012).

50. K. C. Hansen, A. D’Alessandro, C. C. Clement, L. Santambrogio, Lymph formation, composition and circulation: a proteomics perspective. Int Immunol 27, 219–227 (2015).

51. D. M. Heuberger, R. A. Schuepbach, Protease-activated receptors (PARs): mechanisms of action and potential therapeutic modulators in PAR-driven inflammatory diseases. Thromb J 17, 4 (2019).

52. A. Briot et al., Kallikrein 5 induces atopic dermatitis-like lesions through PAR2-mediated thymic stromal lymphopoietin expression in Netherton syndrome. J Exp Med 206, 1135–1147 (2009).

53. J. M. Braz et al., Genetic priming of sensory neurons in mice that overexpress PAR2 enhances allergen responsiveness. Proc Natl Acad Sci U S A 118, (2021).

54. A. Briot et al., Par2 inactivation inhibits early production of TSLP, but not cutaneous inflammation, in Netherton syndrome adult mouse model. J Invest Dermatol 130, 2736–2742 (2010).

55. T. P. Barr et al., PAR2 Pepducin-Based Suppression of Inflammation and Itch in Atopic Dermatitis Models. J Invest Dermatol 139, 412–421 (2019).

56. S. H. Wang, Y. G. Zuo, Thymic Stromal Lymphopoietin in Cutaneous Immune-Mediated Diseases. Front Immunol 12, 698522 (2021).

57. J. M. Leyva-Castillo, P. Hener, H. Jiang, M. Li, TSLP produced by keratinocytes promotes allergen sensitization through skin and thereby triggers atopic march in mice. J Invest Dermatol 133, 154–163 (2013).

58. E. E. West, M. Kashyap, W. J. Leonard, TSLP: A Key Regulator of Asthma Pathogenesis. Drug Discov Today Dis Mech 9, (2012).

59. J. Yoo et al., Spontaneous atopic dermatitis in mice expressing an inducible thymic stromal lymphopoietin transgene specifically in the skin. J Exp Med 202, 541–549 (2005).

60. J. F. Lai, L. J. Thompson, S. F. Ziegler, TSLP drives acute TH2-cell differentiation in lungs. J Allergy Clin Immunol 146, 1406–1418 e1407 (2020).

61. M. D. Howell et al., Th2 cytokines act on S100/A11 to downregulate keratinocyte differentiation. J Invest Dermatol 128, 2248–2258 (2008).

62. S. H. Lee et al., Ameliorating effect of dipotassium glycyrrhizinate on an IL-4- and IL-13-induced atopic dermatitis-like skin-equivalent model. Arch Dermatol Res 311, 131–140 (2019).

63. M. Furue, Regulation of Filaggrin, Loricrin, and Involucrin by IL-4, IL-13, IL-17A, IL-22, AHR, and NRF2: Pathogenic Implications in Atopic Dermatitis. Int J Mol Sci 21, (2020).

64. J. Kobayashi et al., Reciprocal regulation of permeability through a cultured keratinocyte sheet by IFN-gamma and IL-4. Cytokine 28, 186–189 (2004).

65. T. Zheng et al., Transgenic expression of interleukin-13 in the skin induces a pruritic dermatitis and skin remodeling. J Invest Dermatol 129, 742–751 (2009).

66. B. J. Mehrara et al., Pilot Study of Anti-Th2 Immunotherapy for the Treatment of Breast Cancer-Related Upper Extremity Lymphedema. Biology (Basel) 10, (2021).

67. K. H. Hanel, C. Cornelissen, B. Luscher, J. M. Baron, Cytokines and the skin barrier. Int J Mol Sci 14, 6720–6745 (2013).

68. A. Bar-Or, A. Pachner, F. Menguy-Vacheron, J. Kaplan, H. Wiendl, Teriflunomide and its mechanism of action in multiple sclerosis. Drugs 74, 659–674 (2014).

69. A. Jasiecka-Mikolajczyk, P. Socha, Teriflunomide inhibits activation-induced CD25 expression on T cells and may affect Foxp3-expressing regulatory T cells. Res Vet Sci 132, 17–27 (2020).

70. C. K. Derian, A. J. Eckardt, P. Andrade-Gordon, Differential regulation of human keratinocyte growth and differentiation by a novel family of protease-activated receptors. Cell Growth Differ 8, 743–749 (1997).

71. J. F. Hsu, R. P. Yu, E. W. Stanton, J. Wang, A. K. Wong, Current Advancements in Animal Models of Postsurgical Lymphedema: A Systematic Review. Adv Wound Care (New Rochelle) 11, 399–418 (2022).

72. A. A. Grada, T. J. Phillips, Lymphedema: Pathophysiology and clinical manifestations. J Am Acad Dermatol 77, 1009–1020 (2017).

73. G. D. Garcia Nores et al., Regulatory T cells mediate local immunosuppression in lymphedema. J Invest Dermatol 138, 325–335 (2018).

74. I. Martinez-Corral et al., Vegfr3-CreER (T2) mouse, a new genetic tool for targeting the lymphatic system. Angiogenesis 19, 433–445 (2016).

75. S. Z. Aschen et al., Lymph node transplantation results in spontaneous lymphatic reconnection and restoration of lymphatic flow. Plast Reconstr Surg 133, 301–310 (2014).

76. N. W. Clavin et al., TGF-beta1 is a negative regulator of lymphatic regeneration during wound repair. American journal of physiology. Heart and circulatory physiology 2008 Nov;295(5):H2113–27. doi, 10.1152/ajpheart.00879.02008.

