## Supplementary data for "Keratinocytes coordinate inflammatory responses and regulate development of secondary lymphedema"

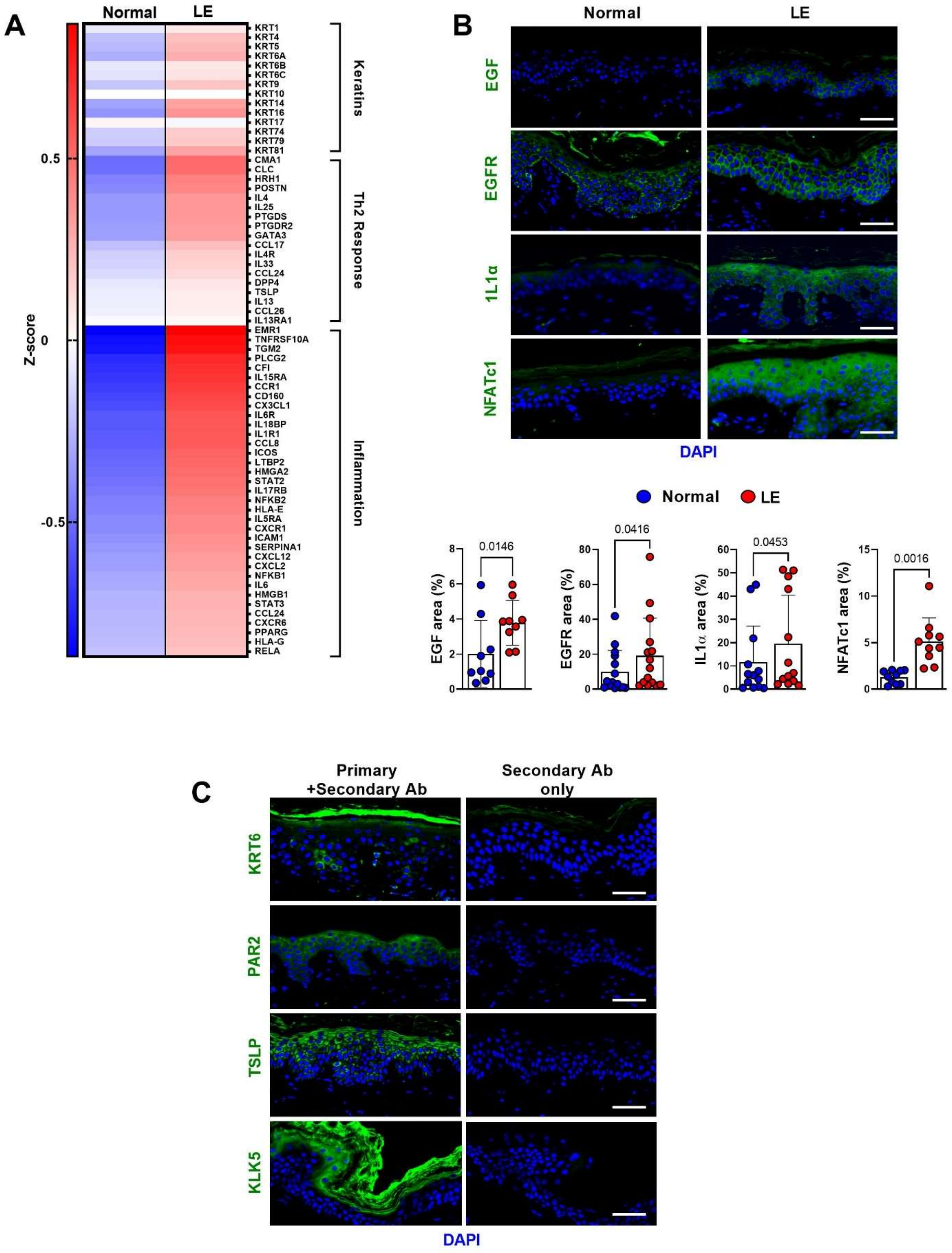

A
Normal
LE
Z-score
0.5
0
-0.5
KRT1
KRT4
KRT5
KRT6A
KRT6B
KRT6C
KRT9
KRT10
KRT14
KRT16
KRT17
KRT74
KRT79
KRT81
Keratins
CMA1
CLC
HRH1
POSTN
IL4
IL25
PTGDS
PTGDR2
GATA3
CCL17
IL4R
IL33
CCL24
DPP4
TSLP
IL13
CCL26
IL13RA1
Th2 Response
EMR1
TNFRSF10A
TGM2
PLCG2
CFI
IL15RA
CCR1
CD160
CX3CL1
IL6R
IL18BP
IL1R1
CCL8
ICOS
LTBP2
HMGA2
STAT2
IL17RB
NFKB2
HLA-E
IL5RA
CXCR1
ICAM1
SERPINA1
CXCL12
CXCL2
NFKB1
IL6
HMGB1
STAT3
CCL24
CXCR6
PPARG
HLA-G
RELA
Inflammation
B
Normal
LE
EGF
EGFR
1L1α
NFATc1
DAPI
Normal
LE
EGF area (%)
0.0146
EGFR area (%)
0.0416
IL1α area (%)
0.0453
NFATc1 area (%)
0.0016
C
Primary +Secondary Ab
Secondary Ab only
KRT6
PAR2
TSLP
KLK5
DAPI

**Figure S1. Expression of keratinocyte growth factors and inflammatory cytokines are upregulated in lymphedema.**

A. Expression of keratins, Th2 response, and inflammatory genes by RNAseq in normal and lymphedematous (LE) skin biopsies from patients with unilateral BCRL (N=4). Each box represents the average mRNA expression of 4 patients.

B. Representative immunofluorescent images (*top*) and quantification (*bottom*) of EGF, EGFR, IL1α, and NFATc1 area in normal and lymphedematous (LE) skin biopsies from patients with unilateral BCRL. Scale bar: 50 μm. Each circle represents the average quantification of 3 HPF views for each patient (N=10-15). *P* values were calculated by paired student's t-test.

C. Immunofluorescent analysis of lymphedema skin biopsies with each antibody (Ab) and respective negative controls without primary antibodies. Scale bar: 50 μm.

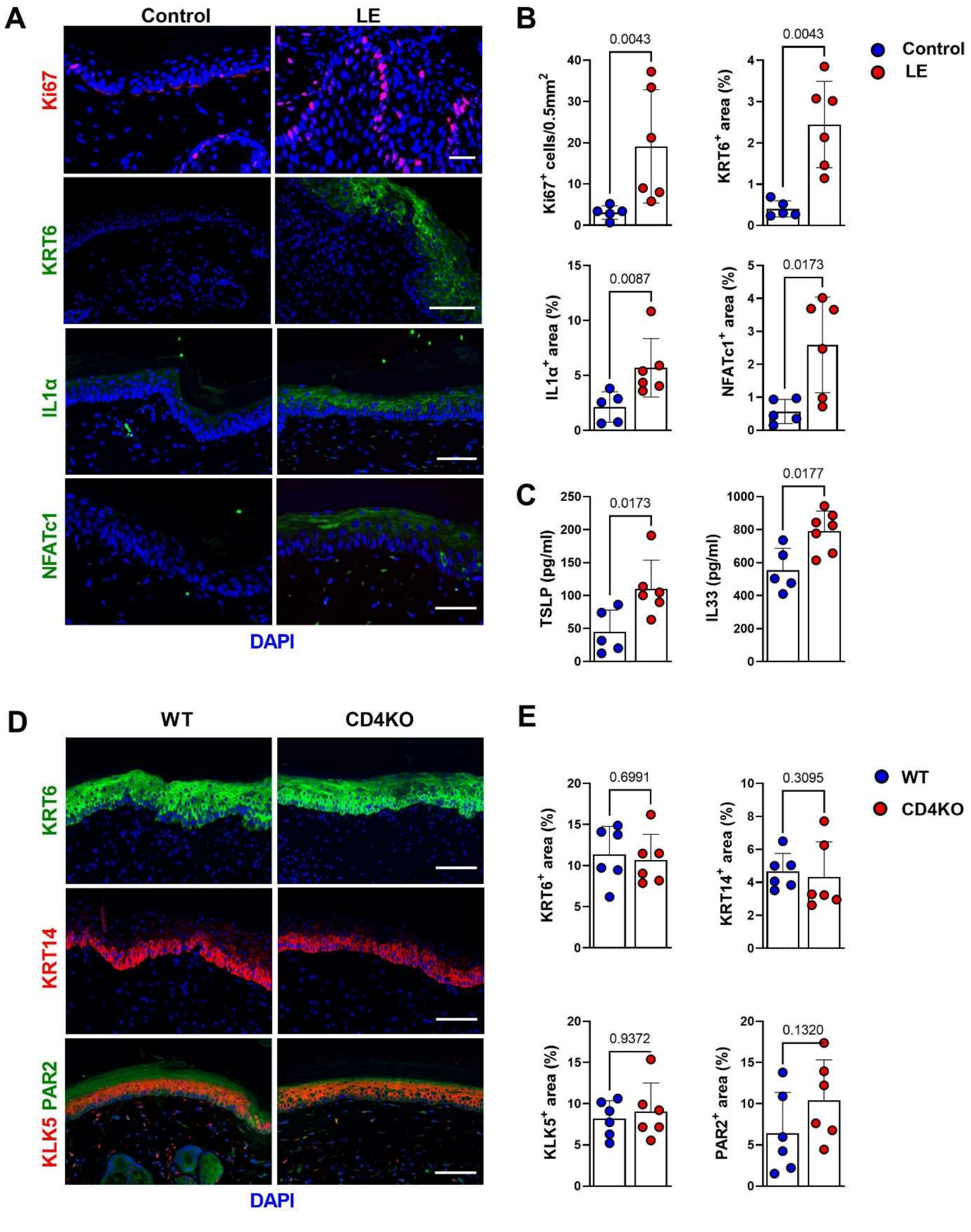

A
Control
LE
Ki67
KRT6
IL1α
NFATc1
DAPI
B
0.0043
Ki67+ cells/0.5mm2
0.0043
KRT6+ area (%)
Control
LE
0.0087
IL1α+ area (%)
0.0173
NFATc1+ area (%)
C
0.0173
TSLP (pg/ml)
0.0177
IL33 (pg/ml)
D
WT
CD4KO
KRT6
KRT14
KLK5 PAR2
DAPI
E
0.6991
KRT6+ area (%)
0.3095
KRT14+ area (%)
WT
CD4KO
0.9372
KLK5+ area (%)
0.1320
PAR2+ area (%)

**Figure S2. Gene expression changes in keratinocytes occur in rapidly after lymphatic injury and are independent of CD4 T cells.**

A. Representative immunofluorescent images of KRT6, Ki67, IL1α, and NFATc1 staining in tail skin harvested 2 weeks after surgery from control and lymphedema (LE) mice. Scale bar: 100 μm.

B. Quantification of KRT6, Ki67, IL1α, and NFATc1 area in tail skin harvested 2 weeks after surgery from control and lymphedema (LE) mice. Each circle represents the average of 3 HPF views of each mouse (N=5-6). *P* values were calculated by Mann-Whitney test.

C. TSLP and IL33 ELISA from protein lysates of tail skin harvested 2 weeks after surgery from control and lymphedema (LE) mice (N=5-7). Each circle represents one mouse. *P* values were calculated by Mann-Whitney test.

D. Representative immunofluorescent images of KRT6, KRT14, KLK5, and PAR2 staining in tail specimens harvested 2 weeks after tail skin and lymphatic excision in wild-type (WT) and CD4 knockout (CD4KO) mice. Scale bar: 100 μm.

E. Quantification of KRT6, KRT14, KLK5, and PAR2 area in tail specimens harvested 2 weeks after tail skin and lymphatic excision in wild-type (WT) and CD4 knockout (CD4KO) mice. Each circle represents the average of 3 HPF views of each mouse (N=6).

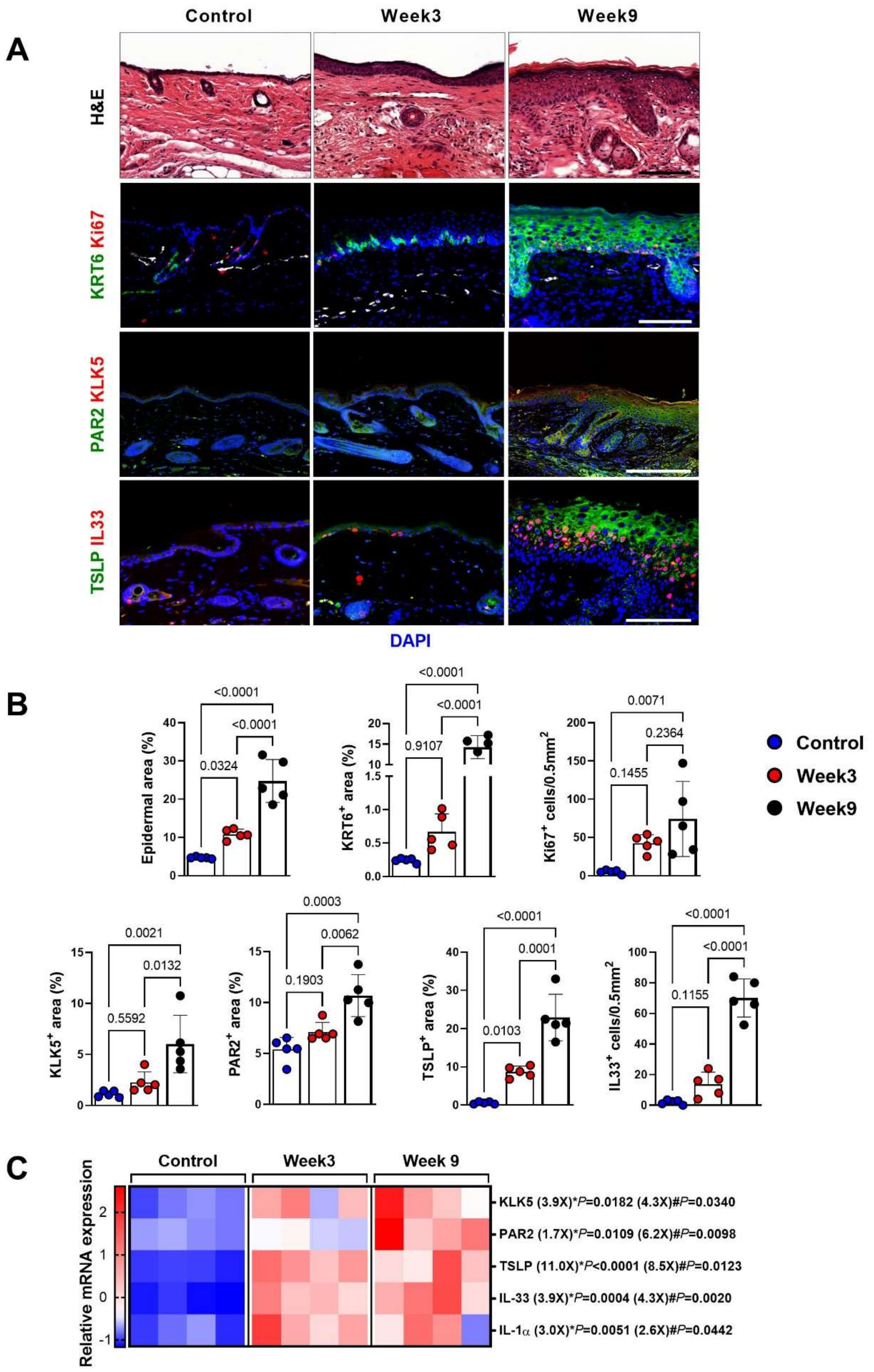

A
Control
Week3
Week9
H&E
KRT6 Ki67
PAR2 KLK5
TSLP IL33
DAPI
B
Epidermal area (%)
<0.0001
<0.0001
0.0324
KRT6+ area (%)
<0.0001
<0.0001
0.9107
Ki67+ cells/0.5mm2
0.0071
0.2364
0.1455
Control
Week3
Week9
KLK5+ area (%)
0.0021
0.0132
0.5592
PAR2+ area (%)
0.0003
0.0062
0.1903
TSLP+ area (%)
<0.0001
0.0001
0.0103
IL33+ cells/0.5mm2
<0.0001
<0.0001
0.1155
C
Control
Week3
Week 9
Relative mRNA expression
KLK5 (3.9X)*P=0.0182 (4.3X)#P=0.0340
PAR2 (1.7X)*P=0.0109 (6.2X)#P=0.0098
TSLP (11.0X)*P<0.0001 (8.5X)#P=0.0123
IL-33 (3.9X)*P=0.0004 (4.3X)#P=0.0020
IL-1α (3.0X)*P=0.0051 (2.6X)#P=0.0442

**Figure S3. Hyperkeratosis and Th2-inducing cytokine expression are increased in a non-surgical model of lymphedema.**

A. Representative H&E and immunofluorescent images of KRT6, Ki67, KLK5, PAR2, TSLP, and IL33 in WT hindlimb skin harvested 9 weeks after DT injection and FLT4CreDTRfloxed hindlimb skin harvested 3 or 9 weeks after DT injection. Scale bar: 100 µm.

B. Quantification of epidermal, KRT6, Ki67, KLK5, PAR2, TSLP, and IL33 area in WT hindlimb skin harvested 9 weeks after DT injection and FLT4CreDTRfloxed hindlimb skin harvested 3 or 9 weeks after DT injection. Each circle represents the average quantification of 3 HPF views for each mouse (N=5). *P* values were calculated by one-way ANOVA.

C. Relative mRNA expression by qPCR in WT hindlimb skin harvested 9 weeks after DT injection and FLT4CreDTRfloxed hindlimb skin harvested 3 or 9 weeks after DT injection. (N=4). mRNA expression was normalized to β-actin expression. Each box represents one mouse. **P* indicates week 3 compared to control; #*P* indicates week 9 compared to control. *P* values were calculated by Mann-Whitney test.

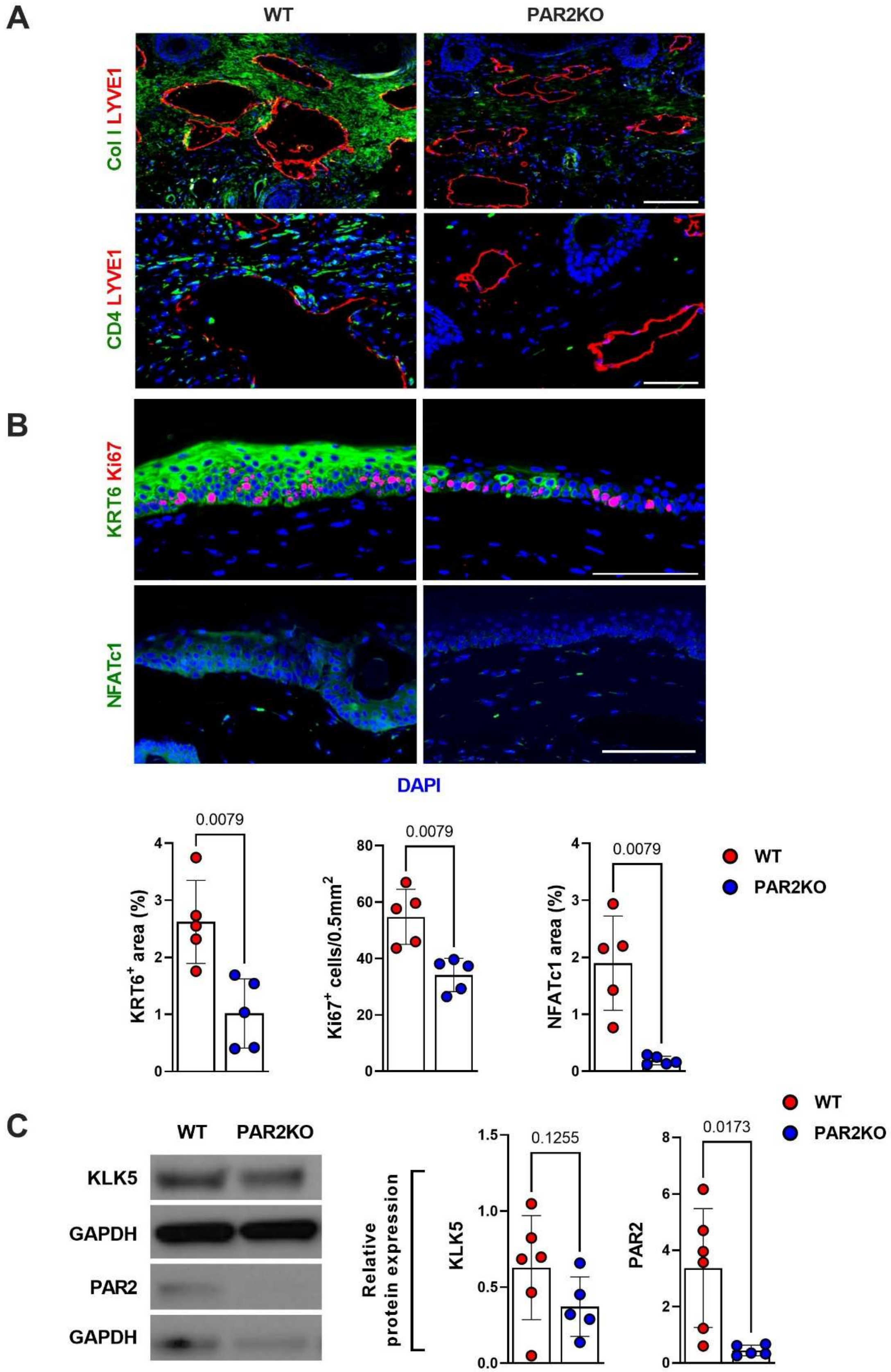
A
WT
PAR2KO
Col I LYVE1
CD4 LYVE1
B
KRT6 Ki67
NFATc1
DAPI
0.0079
KRT6+ area (%)
0.0079
Ki67+ cells/0.5mm2
0.0079
NFATc1 area (%)
WT
PAR2KO
C
WT
PAR2KO
KLK5
GAPDH
PAR2
GAPDH
Relative protein expression
0.1255
KLK5
0.0173
PAR2
WT
PAR2KO

**Figure S4. PAR2 knockout decreases fibrosis, CD4$^+$ cell infiltration, and hyperkeratosis after lymphatic injury**

A. Representative immunofluorescent images of collagen I, LYVE1, and CD4 staining in lymphedema tail skin of wild-type (WT) and PAR2 knockout (PAR2KO) mice harvested 6 weeks after tail skin and lymphatic excision. Scale bar: 100 μm.

B. Representative immunofluorescent images (*top*) and quantification (*bottom*) of KRT6, Ki67, and NFATc1 area in tail skin of WT and PAR2KO mice harvested 6 weeks after tail skin and lymphatic excision. Scale bar: 100 μm. Each circle represents the average quantification of 3 HPF views for each mouse (N=5; *bottom*). *P* values were calculated by Mann-Whitney test.

C. Representative western blots of KLK5 and PAR2 in lymphedema tail skin of WT and PAR2KO mice harvested 6 weeks after tail skin and lymphatic excision (*left*) and quantification relative to GAPHD (*right*). Each circle represents each mouse (N=5-6). *P* values were calculated by Mann-Whitney test.

A

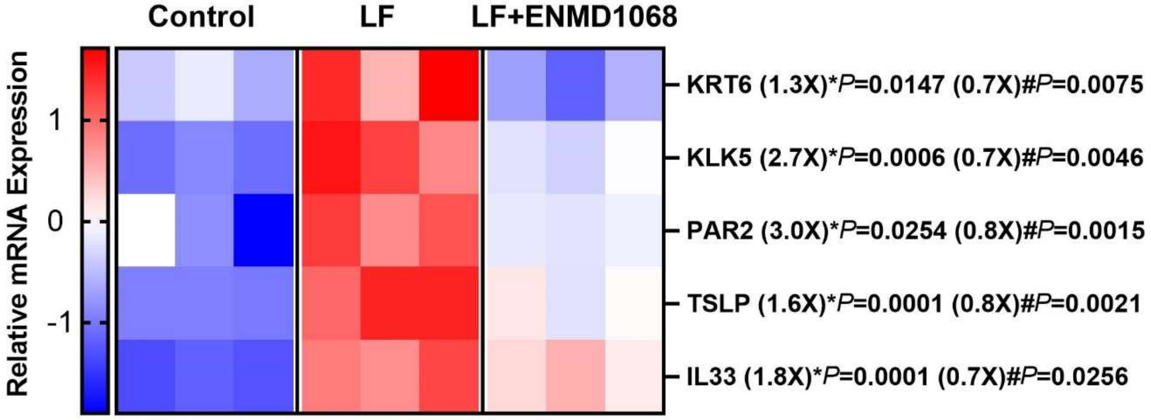

Control
LF
LF+ENMD1068
Relative mRNA Expression
1
0
-1
KRT6 (1.3X)*P=0.0147 (0.7X)#P=0.0075
KLK5 (2.7X)*P=0.0006 (0.7X)#P=0.0046
PAR2 (3.0X)*P=0.0254 (0.8X)#P=0.0015
TSLP (1.6X)*P=0.0001 (0.8X)#P=0.0021
IL33 (1.8X)*P=0.0001 (0.7X)#P=0.0256


B

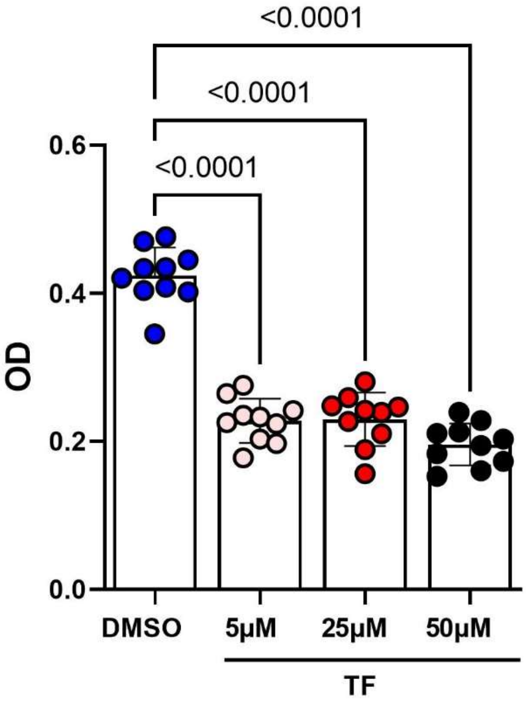

<0.0001
<0.0001
<0.0001
OD
0.6
0.4
0.2
0.0
DMSO
5μM
25μM
50μM
TF

**Figure S5. ENMD1068 and teriflunomide decrease keratinocyte activation in response to lymphatic fluid.**

A. Relative mRNA expression by qPCR in cultured h-keratinocytes treated with PBS (control), lymphatic fluid (LF), or LF+ENMD (N=3). mRNA expression was normalized by β-actin expression. Each box represents each experiment with independently cultured keratinocytes. Fold changes relative to control for LF, and relative to LF for LF+ENMD. **P* indicates LF compared to control, #*P* indicates for LF+ENMD compared to LF. *P* values were calculated by Mann-Whitney test.

B. Proliferation (MTT assay) of h-keratinocytes cultured with DMSO only or teriflunomide (TF) at the indicated concentrations (N=10). Each circle represents an individual experiment. *P* values were calculated by one-way ANOVA.

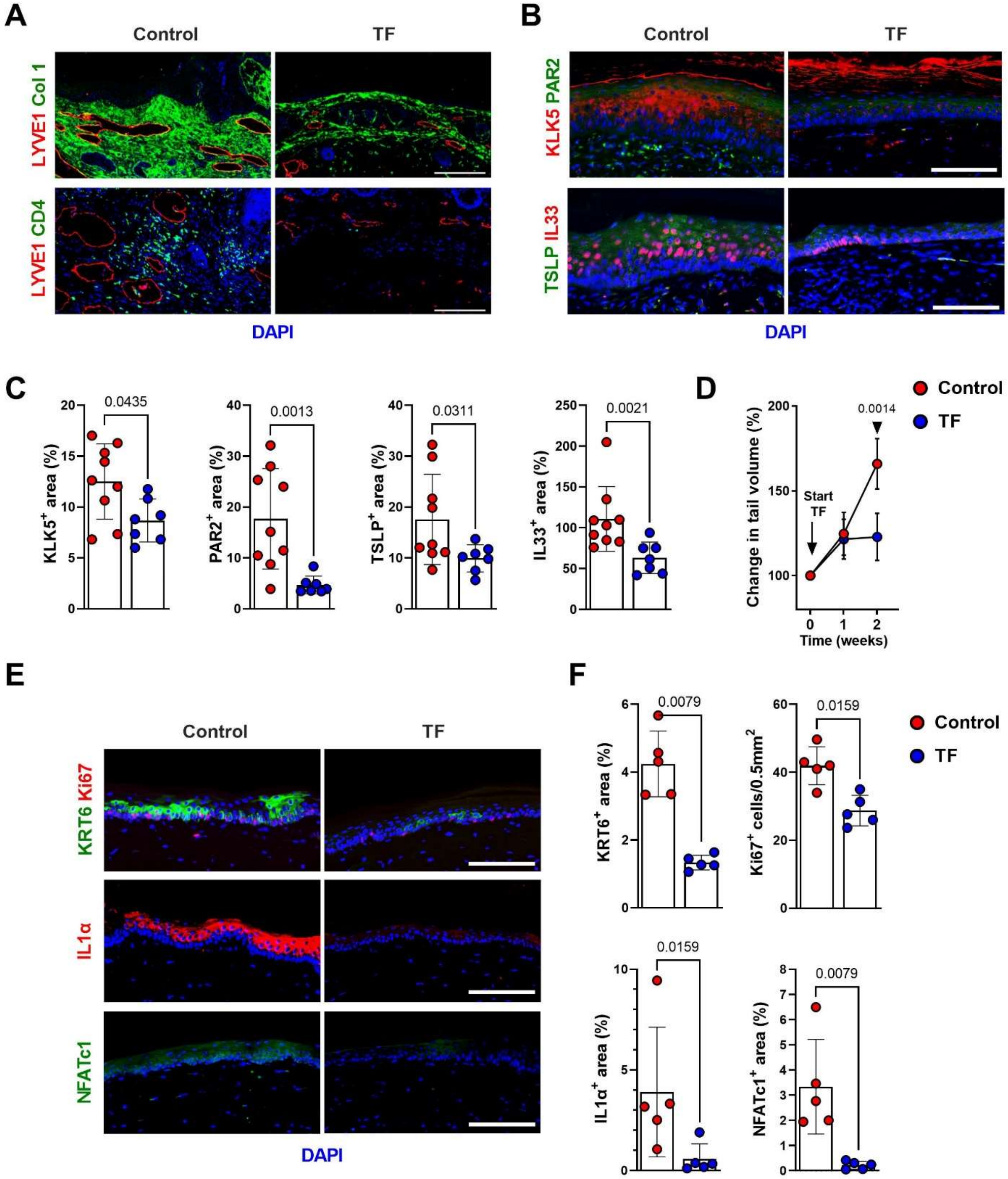
A
Control
TF
LYVE1 Col 1
LYVE1 CD4
DAPI
B
Control
TF
KLK5 PAR2
TSLP IL33
DAPI
C
0.0435
KLK5+ area (%)
0.0013
PAR2+ area (%)
0.0311
TSLP+ area (%)
0.0021
IL33+ area (%)
D
Change in tail volume (%)
Start TF
0.0014
Time (weeks)
Control
TF
E
Control
TF
KRT6 Ki67
IL1α
NFATc1
DAPI
F
0.0079
KRT6+ area (%)
0.0159
Ki67+ cells/0.5mm2
0.0159
IL1α+ area (%)
0.0079
NFATc1+ area (%)
Control
TF

**Figure S6. Teriflunomide treatment decreases fibrosis, $CD4^+$ infiltration, and hyperkeratosis after lymphatic injury.**

A. Representative immunofluorescent images of collagen I, LYVE1, and CD4 staining in tail skin samples harvested from mice treated with vehicle (control) or teriflunomide (TF) once daily for 4 weeks beginning 2 weeks after tail skin and lymphatic excision. Scale bar: 100 µm.

B. Representative immunofluorescent images of KLK5, PAR2, TSLP, and IL33 staining in tail skin samples harvested from mice treated with vehicle (control) or teriflunomide (TF) once daily for 4 weeks beginning 2 weeks after tail skin and lymphatic excision. Scale bar: 100 µm.

C. Quantification of KLK5, PAR2, TSLP, and IL33 area in vehicle and TF-treated mice. Each circle represents the average quantification of 3 HPF views for each mouse (N=7-9). *P* values were calculated by Mann-Whitney test.

D. Changes in tail volume over time in mice treated with vehicle (control) or TF for 2 weeks starting one day after tail skin and lymphatic excision. Each circle represents the average measurement from each mouse (N=5). *P* values were calculated by 2-way ANOVA with multiple comparisons.

E. Representative immunofluorescent images of KRT6, Ki67, IL1α, and NFATc1 staining. of tail skin from mice treated with vehicle (control) or TF for 2 weeks starting one day after tail skin and lymphatic excision. Scale bar: 100 µm.

F. Quantification of KRT6, Ki67, IL1α, and NFATc1 area in vehicle and TF-treated mice. Each circle represents the average quantification of 3 HPF views for each mouse (N=5). *P* values were calculated by Mann-Whitney test.

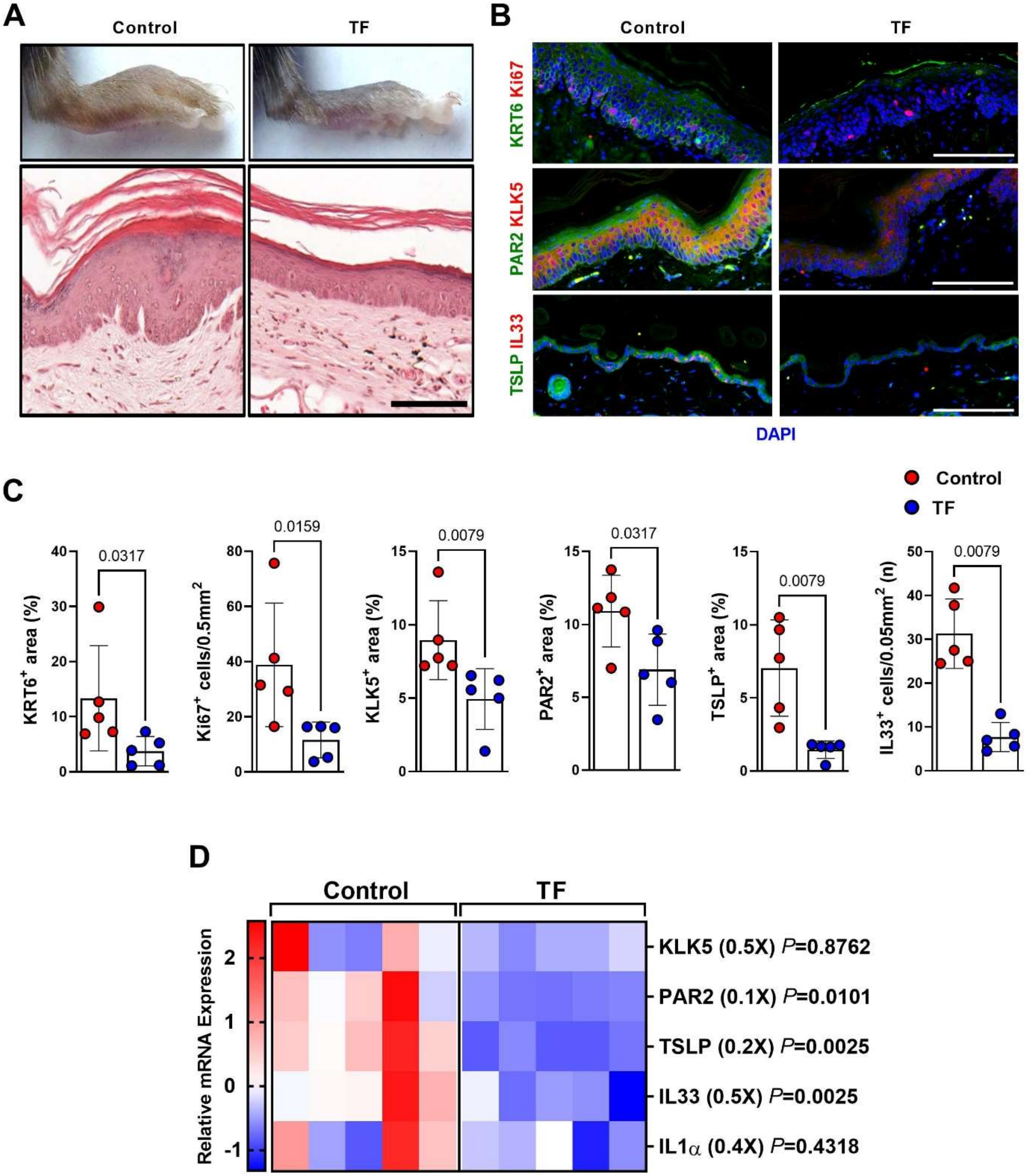
A
Control
TF
B
Control
TF
KRT6 Ki67
PAR2 KLK5
TSLP IL33
DAPI
C
Control
TF
0.0317
0.0159
0.0079
0.0317
0.0079
0.0079
KRT6+ area (%)
Ki67+ cells/0.5mm2
KLK5+ area (%)
PAR2+ area (%)
TSLP+ area (%)
IL33+ cells/0.05mm2 (n)
D
Control
TF
Relative mRNA Expression
KLK5 (0.5X) P=0.8762
PAR2 (0.1X) P=0.0101
TSLP (0.2X) P=0.0025
IL33 (0.5X) P=0.0025
IL1α (0.4X) P=0.4318

**Figure S7. Teriflunomide decreases lymphedema in a non-surgical mouse model.**

A. Representative images of lymphedema hindlimb and H&E images of lymphedema hindlimb skin from mice treated with vehicle (control) or teriflunomide (TF) for 8 weeks starting a week after DT injection. Scale bar: 100 μm.

B. Representative immunofluorescent images of KRT6, Ki67, KLK5, PAR2, TSLP, and IL33 staining of hindlimb skin from control and TF-treated mice. Scale bar: 100 μm.

C. Quantification of KRT6, Ki67, KLK5, PAR2, TSLP, and IL33 area in hindlimb skin from control and TF-treated mice. Each circle represents the average quantification of 3 HPF views for each mouse (N=5). *P* values were calculated by Mann-Whitney test.

D. Relative mRNA expression by qPCR in hindlimb skin from vehicle (control) and TF-treated mice (N=5). mRNA expression was normalized to β-actin expression. Each box represents one mouse. *P* values were calculated by Mann-Whitney test. Fold change from control is shown in parentheses.
